# A machine vision based frailty index for mice

**DOI:** 10.1101/2021.09.27.462066

**Authors:** Leinani E. Hession, Gautam S. Sabnis, Gary A. Churchill, Vivek Kumar

## Abstract

Chronological aging is uniform, but biological aging is heterogeneous. Clinically, this heterogeneity manifests itself in health status and mortality, and it distinguishes healthy from unhealthy aging. Frailty indexes (FIs) serve as an important tool in gerontology to capture health status. FIs have been adapted for use in mice and are an effective predictor of mortality risk. To accelerate our understanding of biological aging, high-throughput approaches to pre-clinical studies are necessary. Currently, however, mouse frailty indexing is manual and relies on trained scorers, which imposes limits on scalability and reliability. Here, we introduce a machine learning based visual frailty index (vFI) for mice that operates on video data from an open field assay. We generate a large mouse FI dataset of both male and female mice. From video data on these same mice, we use neural networks to extract morphometric, gait, and other behavioral features that correlate with manual FI score and age. We use these features to train a regression model that accurately predicts the normalized FI score within 0.04 ± 0.002 (mean absolute error). We show that features of biological aging are encoded in open-field video data and can be used to construct a vFI that can complement or replace current manual FI methods. We use the vFI data to examine sex-specific aspects of aging in mice. This vFI provides increased accuracy, reproducibility, and scalability, that will enable large scale mechanistic and interventional studies of aging in mice.

## 2 Introduction

Aging is a terminal process that affects all biological systems. Biological aging – in contrast to chronological aging – occurs at different rates for different individuals. In humans, growing old comes with increased health issues and mortality rates, yet some individuals live long and healthy lives while others succumb earlier to diseases and disorders. More precisely, there is an observed heterogeneity in mortality risk and health status among individuals within an age cohort [1, 2]. The concept of frailty is used to quantify this heterogeneity and is defined as the state of increased vulnerability to adverse health outcomes [3]. Identifying frailty is clinically important as frail individuals have increased risk of diseases and disorders, worse health outcomes from the same disease, and even different symptoms of the same disease [2].

The frailty index (FI) is an invaluable and widely used tool which outperforms other methods to quantify frailty [4]. In this method, an individual is scored on a set of age-related health deficits to produce a cumulative score. Each deficit must have the following characteristics: they must be health related, they must increase in the population with age, and they must not saturate in the population too early [5]. The presence and severity of each health deficit is scored as 0 for not present, 0.5 for partially present, or 1 for present. A compelling finding of FIs is that the exact health deficits scored can vary between indexes but still show similar characteristics and utility [5]. That is, two sufficiently large FIs with different numbers and selections of deficits scored would still show a similar average rate of deficit accumulation with age and the same submaximal limit (the highest FI score observed). More importantly, both FIs would predict an individual’s risk of adverse health outcomes, hospitalization, and mortality. This feature of FIs is advantageous as researchers can pull data from varied large health databases, aiding in large-scale studies. It also suggests that frailty is a legitimate phenomenon and that FIs are a valid measure of biological aging. Different people age not only at different rates but in different ways; one person may have severe mobility issues but have a sharp memory, while another may have a healthy heart but a weak immune system. Both may be equally frail, but this is only made clear by sampling a variety of health deficits. Indeed, FI scores outperform other developed measures including molecular markers and frailty phenotyping at predicting mortality risk and health status [4, 6, 7].

FIs have been adapted for use in mice using a variety of both behavioral and physiological measures as index items [2, 4, 8]. Mouse FI shows many of the characteristics of human FIs, including a submaximal limit and a strong correlation with mortality [9]. Mouse FIs have been successfully used to evaluate a variety of aging interventions [10] and for construction of models of chronological age and mortality [4]. Unlike human FIs, the majority of mouse frailty indexing has been performed using the same set of health-deficits as Whitehead et al. [2], although some studies have substituted a small number of mouse FI items for ones that better fit their specific strain or experiment [10].

The successful creation of the mouse FI is a major step forward in aging research, particularly for interventional studies that follow health outcomes over long periods of time and are often carried out by multiple labs or at disparate locations. However, because conducting mouse FI requires trained individuals for manual scoring, it often limits the scalability of the tool; FI scoring of hundreds of mice is a labor-intensive task. While human studies can draw from large health databases and have sample sizes in the thousands, mouse studies that employ FIs are much smaller due to the low-throughput nature of the scoring [9]. Furthermore, since many of the FI metrics require some level of subjective judgment, there are concerns about scorer-based variability and reproducibility [10–12]. Reliability of the FI between scorers has been found to be very good, although recent work has shown that the professional background of scorers affects FI scores; specifically, inter-scorer reproducibility was poor between animal technicians and research scientists [12]. These studies on inter-scorer agreement strongly emphasize the importance of inter-scorer discussion and refinement in obtaining high agreement [10–12]. In fact, it has been shown that in the absence of discussion and refinement, inter-rater reliability does not improve solely by practice and experience [12]. This discussion and refinement is not always feasible in multi-site or long-term studies. Therefore, although the FI is an extremely useful tool for aging research, an increase in its scalability, reliability, and reproducibility through automation would enhance its utility.

Towards this end, we developed an automated visual FI using video of mice in the open field. The open field is one of the oldest and most widely used assays for rodent behavior [13]. Commonly, measures like total distance travelled, thigmotaxis, grooming bouts, urination, and defecation have been used to infer the behavioral state of the animal [14]. However, advances in machine learning techniques have greatly expanded the types of metrics that can be extracted from the open field assay beyond the traditional metrics of hyperactivity and anxiety [15, 16]. These advances are largely due to discoveries in the computer vision and statistical learning field [17–22]. We along with a number of other groups have applied these new methods to animal behavior analysis. Our group has developed methods for image segmentation and tracking in complex environments [23], action detection [24] and pose-based gait and whole body coordination measurements in the open field [25]. These and other highly sensitive methods have advanced animal behavior extraction [15, 26–28].

Our goal was to develop an efficient scalable method to determine frailty in the mouse using computer-vision based features. We hypothesized that biological aging produces changes in behavior and physiology that are encoded in video data, i.e. we can visually determine the frailty of an animal based on their open field behavior. Additionally, sex differences in the FI are still an active area of research in mice [9, 29]. Many age related changes are known to be sex-specific in humans [30–32]. Therefore, we generated one of the largest mouse FI datasets consisting of both males and females. We extracted sensitive measures of mouse movement (gait and posture), morphometric, and other behaviors using machine learning methods. We used these features to construct a vFI assay that has high prediction accuracy. Through modeling we also gained insight into which video features are important to predict FI score across age and frailty status. Our automated vFI will increase efficiency and accuracy for large scale studies that explore mechanisms and interventions of aging.

## 3 Results

### 3.1 Data collection and study design

Our overall approach is described in Figure 1A. The study was conducted with 643 data points (371 males, 272 females) taken over three rounds of testing with 533 unique mice. The first round (batch 1) of testing included 222 mice (141 males, 81 females). The second round (batch 2) of testing occurred about 5 months later, and included 319 mice (173 males, 146 females). Of those mice, 105 were repeated from the first batch. The third round (batch 3) of testing occurred about a year later in response to reviewer comments with 102 mice (57 males, 45 females). Of those mice, 18 had been previously tested in the first round and 15 had been tested in the second round (Table S1). Top-down video of each mouse in a 1-hour open field session was collected according to previously published protocols [16, 23] (see Methods, Figure 1A, and supplementary Video 1 and 2 as examples of a young and old mouse). Following the open field, each mouse was scored using a standard mouse frailty indexing by a trained expert from the Nathan Shock Center for Aging to assign a manual FI score [33] (Figure S2A). Due to the slight bimodality of our data, we tested for the presence of Simpson’s Paradox [34] and did not see any evidence (Figure S4). The open field video was processed by a tracking network and a pose estimation network, to produce a track, an ellipse-fit, and a 12-point pose of the mouse for each frame [23, 25]. These frame-by-frame measurements were used to calculate a variety of per-video features, including traditional open field measures of anxiety and hyperactivity [23], grooming [24], gait and posture measures [25], and engineered features. All features are defined in Section 3.2 and Table S2 and were used to train a machine learning model to predict chronological and biological age, a visual FI (vFI).

**Figure 1:**
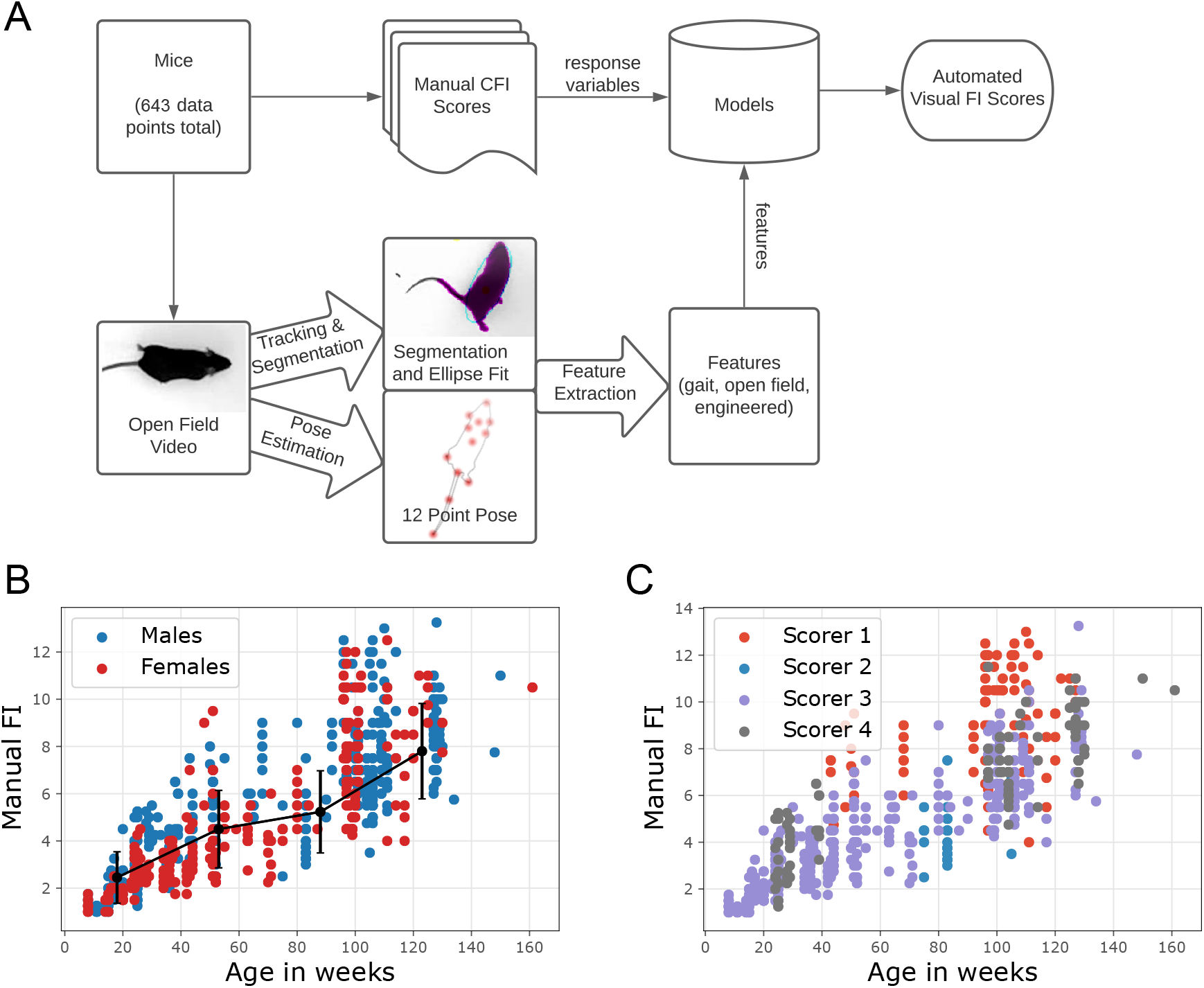
Approach overview to build a visual frailty index. (A) Pipeline for automated visual frailty index (vFI). Top-down videos of the open field for each mouse are processed by a tracking and segmentation network and a pose estimation network. The resulting frame-by-frame ellipse-fits and 12 point pose coordinates are further processed to produce per-video metrics for the mouse. The mouse is also manually frailty indexed to produce a FI score. The video features for each mouse are used to model its FI score. B) Distribution of FI score by age. The black line shows a piece-wise linear fit to the data and the error bars are the standard deviations. C) Effect of scorer on FI data.

Consistent with previous data, in our dataset, mean FI score increases with age (Figure 1B). Heterogeneity of FI scores (shown by the standard deviation bars) also increases with age. We find a sub-maximal limit of FI score slightly below 0.5 for our data, which falls within a range of submaximal limits shown in mice [2, 9]. These results show that our FI data are typical of other mouse data and mirrors the characteristics of human FIs with an increase in average FI scores and heterogeneity of FI scores with age [9]. Over the course of the data collection, 4 different scorers conducted the manual FI. Visual inspection of the data showed a scorer effect on the manual FI score (Figure 1C). For instance, Scorer 1 and 2 tended to generate high and low frailty scores, respectively (Figure S1 B). Modeling indicated that 42% of the variability in manual FI scores was due to a scorer effect (RLRT = 183.85, p *<* 2.2*e*^−16^). A closer examination of which FI items are most affected by scorer showed that piloerection, kyphosis, vision are the most subjective (Figure S1 A). This analysis suggests that scorer effect is an important source of variability in mouse clinical FI.

### 3.2 Feature extraction

The frame-by-frame segmentation, ellipse fit, and 12-point pose coordinates were used to extract pervideo features [23–25]. All extracted features with explanation and source of the measurements can be found in Table S2. Overall, there was a very high correlation between median and mean video metrics (Figure S3A,B). We decided to use only medians and inter-quartile ranges in modeling where possible for two reasons: medians tend to have higher correlation with FI score than means, and medians and inter-quartile ranges are more robust to outlier effects than means and standard deviations, respectively. This gave us a total of 44 video features (Table S3, Table S2). We first looked at metrics taken in standard open field assays such as total locomotor activity, time spent in the periphery vs. center, and grooming bouts (Figure 2A). All standard open-field measures showed low correlation with both FI score and age (Table S3 and SS4).

**Figure 2:**
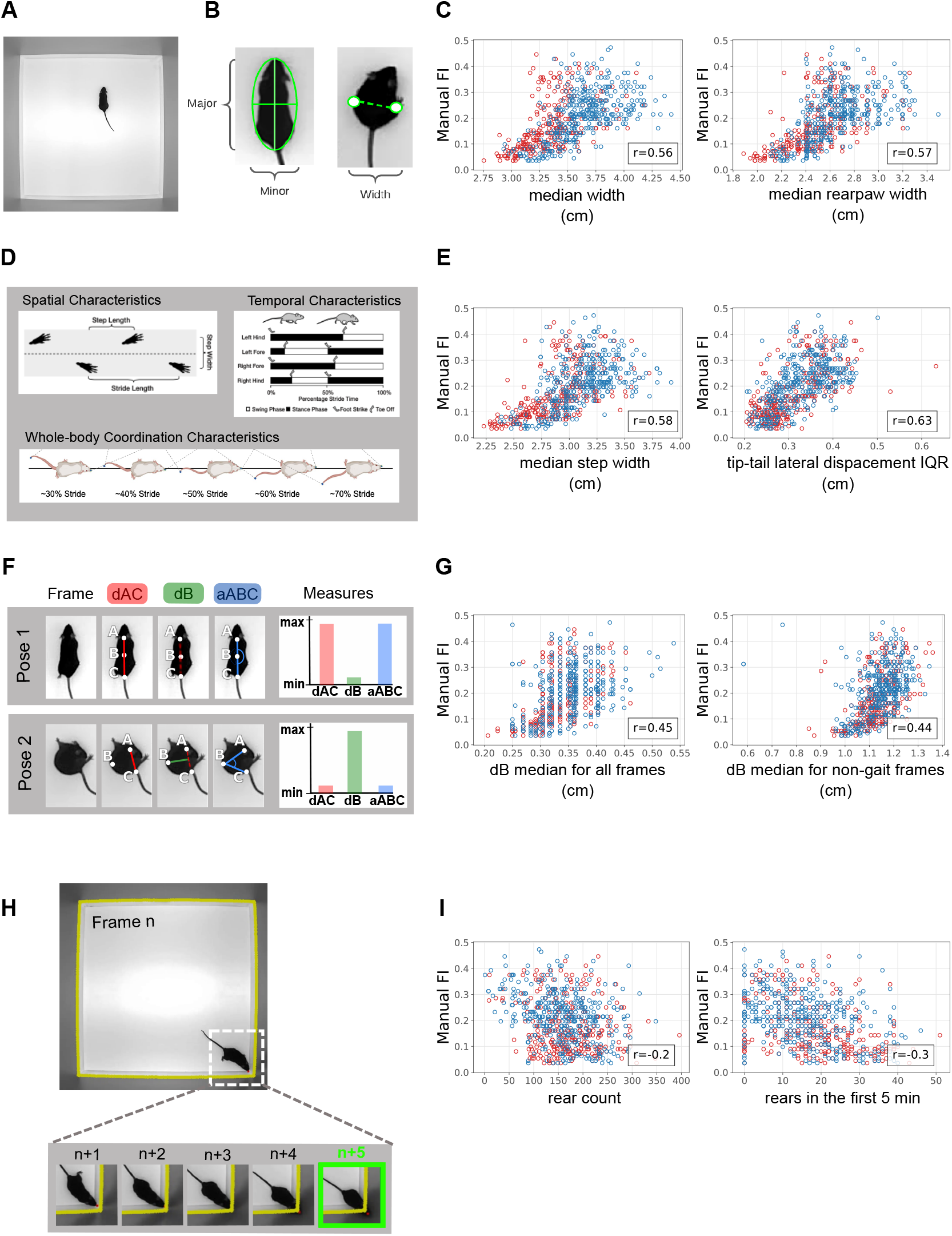
Sample features used in the vFI (A) Single frame of the top-down open-field video. (B) Morphopemetric features from ellipse-fit and rear-paw distance measure performed on the mouse frame by frame. The major and minor axis of the ellipse fit are taken as the length and width respectively. (C) The median ellipse-fit width and the median rear-paw distance taken over all mouse frames are highly correlated with FI score. (D) Spatial, temporal, and whole-body coordination characteristics of gait used to create metrics ([25]). (E) The median step width and the inter-quartile range of tip-tail lateral displacement taken over all strides for a mouse are highly correlated with FI score. (F) Spinal mobility measurements taken at each frame. dAC is the distance between point A and C (base of head and base of tail respectively) normalized for body length, dB is the distance of point B (mid-back) from the midpoint of the line AC, and aABC is the angle formed by the points A, B, and C. When the mouse spine is straight, dAC and aABC are at their maximum value while dB is at its minimum. When the mouse spine is bent, dB is at its maximum value while dAC and aABC are at their minimum. See Supplementary Video 3.(G) The median of dB taken over all mouse frames and the median dB taken only over frames where the mouse is not in gait shows correlation with FI score. (H) Wall rearing event. The contour of the walls of the open field are taken and a buffer of 5 pixels is added (yellow line), marking a threshold. The nose point of the mouse is tracked at each frame. A wall rearing event is defined by the nose point fully crossing the wall threshold. See Supplementary Video 4. (I) The total count of rearing events and the number of rears in the first 5 minutes of the open-field video show some correlation with FI score.

In addition to the existing features, we designed a set of features that we hypothesized may correlate with FI. These include morphometric features that capture the animals shape and size, as well as behavioral features that are associated with flexibility and vertical movement. Changes in body composition and fat distribution with age are observed in humans and rodents [35]. We hypothesized that body composition measurements may show some signal of aging and frailty. We took the major and minor axes of the ellipse fitted to the mouse at each frame as an estimated length and width of the mouse respectively (Figure 2B). The distance between the rear paw coordinates for each frame were taken as another width measurement closer to the hips. The means and medians of the ellipse width, ellipse length, and rear paw width over all frames were used as per-video metrics. Many of these morphometric features showed high correlations with FI score and age (Table S3 and S4), for example, specifically median width and median rear paw width had correlations of r = 0.56 and 0.57, respectively (Figure 2C).

Changes in gait are a hallmark of aging in humans [36, 37] and mice [38, 39]. Recently, we established methods to extract gait and posture measures from freely moving mice in the open field [25]. We carried out similar analysis to explore age-related gait changes in the current cohort of mice (Figure 2D, E). Each stride was analyzed for its spatial, temporal, and posture measures (Figure 2D), resulting in an array of measures of which the medians over all strides for each mouse are taken. We also looked into intra-mouse heterogeneity of gait features using standard deviations and inter-quartile range over all strides for each mouse. Many of these calculated metrics showed a high correlation with FI score and age (Table S3 and SS4), for example, median step width and tip-tail lateral displacement interquartile range (r=0.58 and r=0.63, respectively) (Figure 2E).

We next investigated the bend of the spine throughout the video (see Supplementary Video 3). We hypothesized that aged mice may bend their spine to a lesser degree, or less often due to reduced flexibility or spine mobility. This change in flexibility can be captured by the pose estimation coordinates of three points on the mouse at each video frame: the back of the head (A), the middle of the back (B), and the base of the tail (C). At each frame, the distance between points A and C normalized for mouse length (dAC), the orthogonal distance of the middle of the back B from the line (dB), and the angle of the three points (aABC) were calculated (Figure 2F). For each of the three per-frame measures (dAC, dB, and aABC) a mean, median, standard deviation, minimum, and maximum are calculated per video for all frames and for non-gait frames (frames where the mouse is not in stride). We founnd some moderately high correlations showing relationships between spinal bend and FI score which contradicted our hypothesis (Table S3 and SS4); while we expected dB median for all frames and dB median for non-gait frames to decrease with age, we found that they increase (r=0.45 and 0.44, respectively) (Figure 2G). One possible reason for this result was that very frail mice were spending more time grooming. However, neither grooming bouts or time spent grooming show a strong relationship with FI score (Table S3) or dB median. Another possibility was that high frailty mice walked less and spent more time curled up. This does not seem to be the case either, as there is almost no relationship between either stride count or distance travelled and FI score or dB median. High frailty mice may also have higher dB medians due to body composition, as dB median has a correlation of 0.496 with body weight. It is important to note that these bend metrics cast a wide net; they are an inexpensive and general account of all the activity of the spine during the one-hour open field. Thus, these measures may capture the interaction between body composition and behavior.

While the previous spinal flexibility measures only looked at lateral spinal flexibility, vertical flexibility may also have a relationship to frailty. To investigate this we looked at occurrences of rearing supported by the wall (Figure 2H, Supplementary Video 4). We hypothesized that frailer mice may rear less due to reduced lateral spinal mobility and/or reduced exploratory activity. The edges of the open field were taken and a buffer of 5 pixels added as a boundary. We took the frames in which the nose coordinates of the mouse cross that boundary as instances of rearing. From these heuristic rules we were able to determine the number of rears and the average length of each rearing bout (Table S2). We found that some metrics related to rearing bouts show signal for frailty, specifically total count of rears and rears in the first 5 minutes (r = 0.2 and 0.3, respectively, Figure 2I).

Interestingly, most of the correlations with age were slightly higher than correlations with FI score (Table S3 and SS4). This may be because mice become frail in different ways; one mouse can have high frailty but no dysfunction in their stride width, while on average older mice, regardless of their frailty, have more stride width changes. Of further note is the increase of heterogeneity in many of these measures with both age and FI score (for example: median width, median step width, dB median).

### 3.3 Sex differences in frailty

The sex-specific characteristics of aging are important considerations. To visualize sex differences in frailty, we stratified the FI score data into four age groups and compared the boxplots for each age group between males and females (Figure 3A). The oldest age group included only 9 females compared to 81 males. The range of females frailty scores for each age group tended to fall lower than males except for the oldest age group. The middle two age groups showed highly significant differences in distribution between males and females.

**Figure 3:**
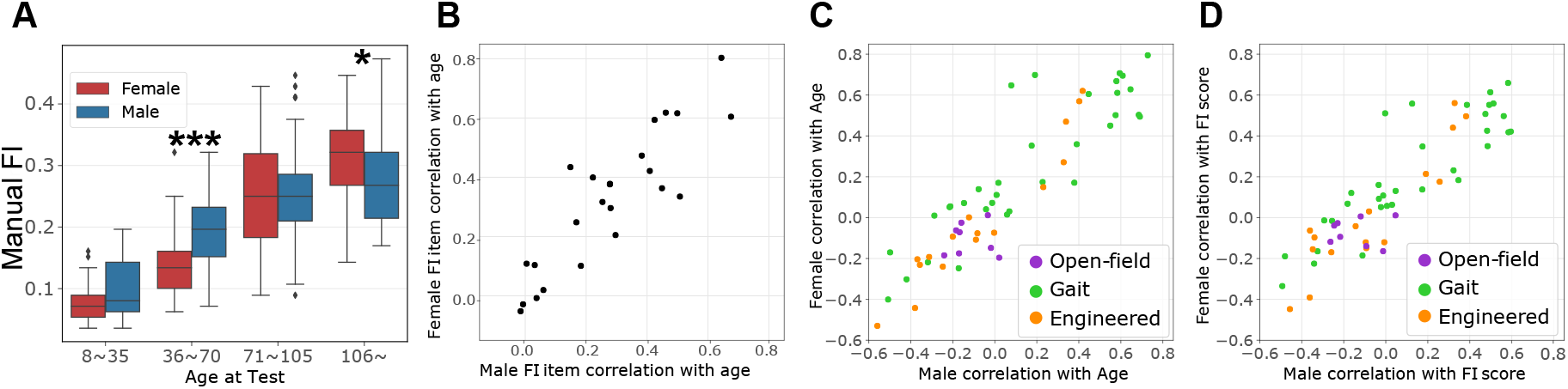
Comparison of male and female FI metrics. (A) The distribution of FI scores for males and females when the data are split into 4 age groups of equal range. The x-ticks represent the midpoint of each age group range. Significant difference in the distribution of male and female scores for that age group determined by the Mann-Whitney U test. (B) Pearson correlations of FI items with age for males compared to females. (C) Pearson correlations of video metrics with FI score for males compared to females. (D) Pearson correlations of video metrics with age for males compared to females.

Comparisons between the correlations of male and female FI item scores with age (Figure 3B), showed an overall high correlation (r=0.85). The average difference between male and female correlations of FI index items with age was 0.08, but a few index items showed notable differences. Alopecia and Menace Reflex have the highest sex differences in their correlation to age (0.29 and 0.21 respectively), with females having a higher correlation for Alopecia and males having a higher correlation for Menace Reflex (TableS5).

The correlations of male and female video features with both FI score and age were also high (r=0.88 and r=0.90 respectively), with an average difference between male and female correlations of video metrics with FI score and age of 0.14 and 0.13 respectively (Table S3 and SS4). In both FI score and age, the video features with the highest sex differences were gait measures related to stride and step length, base tail lateral displacement, and tip tail lateral displacement. The highest sex differences were the correlations between median base tail lateral displacement to age (difference of 0.57) and median tip tail lateral displacement to age (0.50), for which females tended to have a higher correlation to both FI score and age. For the metrics related to stride length and step length (median stride length, difference of 0.33), males had a higher correlation to FI score and age than females. These results show that with age, females significantly increase their base-tail and tip-tail lateral displacement in gait while males show little change in this feature, whereas males show a greater reduction in stride length with age compared to females.

### 3.4 Prediction of age and frailty index from video data

Once we established that our video features correlate with aging and frailty, we used these features as covariates in a model to predict age and manual FI scores (Figure 4A, model vFRIGHT and vFI, respectively). Age is an empirical ground truth and has a strong relationship to frailty. We compared the prediction of age using video features (Figure 4A, Model vFRIGHT) to the prediction of age using manual FI items– a method referred to as the FRIGHT age clock [4] (Figure 4A, Model FRIGHT). We first tested four models - penalized linear regression (LR*) [40], support vector machine (SVM) [41], random forest (RF) [42], and extreme gradient boosting (XGB) [43] (Figure 4B, Panel 1). We selected the random forest regression model to predict age on unseen future data due to its superior performance over other models with a lowest mean absolute error (MAE) (*p <* 2.2*e*^−16^, F_3,147_ = 190.43), root-mean-squared error (RMSE) (*p <* 2.2*e*^−16^, F_3,147_ = 59.53), and highest R^2^ (*p <* 2.2*e*^−16^, F_3,147_ = 58.14) when compared using repeated-measures ANOVA (Figure 4B, Panel 1, Figure S2C). Our vFRIGHT model was able to more accurately and precisely predict age than the FRIGHT clock. vFRIGHT had a superior performance (*p <* 4.7*e* − 5, *F*_1,49_ = 19.9, using repeated-measures ANOVA) with a lower MAE (13.1 ± 0.99 weeks) compared to the FRIGHT clock using FI items (15.7 ± 4 weeks) (Figure 4B, Panel 2). We also compared RMSE (RMSE_vFRIGHT_ = 17.97 ± 1.44, RMSE_FRIGHT_ = 20.62 ± 4.78, *p <* 6.1*e* − 7, *F*_1,49_ = 32.84) and R^2^ (RMSE_vFRIGHT_ = 0.78 ± 0.04, RMSE_FRIGHT_ = 0.76±0.07, *p <* 2.1*e*−8, *F*_1,49_ = 44.54) and found similar significant improvements in predicting age when using video features (Figure S2D). The variance of prediction errors was noticeably reduced for the video based age prediction (vFRIGHT) than the manual FI item based age prediction (FRIGHT) (Figure 4B, Panel 2). These results show that the automated video features more precise information about aging beyond what is addressed in the manual FI items. The video features may also provide information of aging which overlap with the health deficits scored in the manual FI.

**Figure 4:**
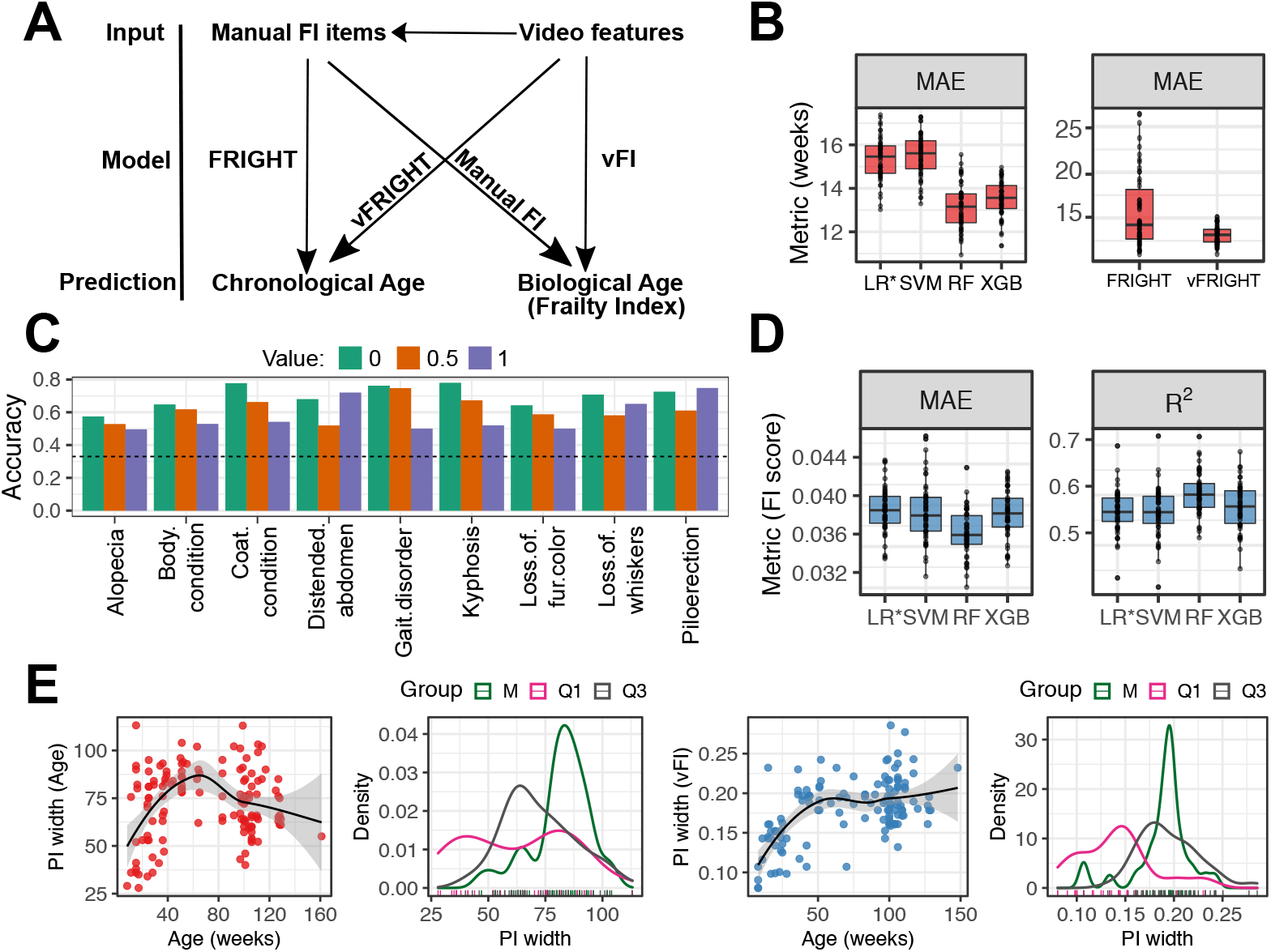
Prediction of age and frailty from video features. (A) A graphical illustration shows different models we fit. (B) Video features are more accurate at predicting age than clinical frailty index items. We compare the performances of random forest models using frailty parameters (FRIGHT) and video-generated features (vFRIGHT) in predicting Age. (C) We reported the performance of our ordinal regression models (classifiers) in terms of accuracy (accurately predicting the value of the frailty parameter in the test using the model trained on the training data). The black dotted line superimposed on the plot shows the accuracy that one would obtain if one guessed the values instead of using the video features. We found that the video features encode helpful information that improves the models’ ability to predict frailty parameters’ values accurately. (D) Comparison among four models (LR*, SVM, RF, XGB) that predict FI score from video features in terms of mean absolute error (MAE) and *R*^2^ shows that RF outperforms other models. (E) We calculate the uncertainty in predicting Age (red) and FI score (blue) and plot it as a function of age (weeks). The black curve shows the loess fit. These plots show less uncertainty in predicting Age and FI scores for very young/old mice. We plot the distributions of prediction interval (PI) widths and find that the PI widths for predicting Age are wider (increased uncertainty in predictions) for mice belonging to the middle age group (M). Similarly, the PI widths for predicting FI scores increase with Age in our data.

To address this, we predicted individual FI items using video features (Figure 4A). Of the 27 items, many had no to almost no non-zero scores, which shows that in our genetically homogeneous data set at least, most of the information in the manual FI are coming from a subset of index items (Figure S2F). We selected only index items with a balanced ratio of 0 to 0.5 and 1 scores for prediction (Figure 4C). We then built a classifier for each of the 9 index items to predict the score given a mouses video features. We predicted the individual FI items’ scores using an ordinal elastic net regression model. For all 9, we were able to predict the score at an accuracy above what would be expected by randomly guessing (Figure 4C, dotted line is guessing accuracy). Many of these FI items have implicit relationships to video features like grooming (ex: coat condition, alopecia), gait/mobility (ex: gait disorders, kyphosis), and body composition (ex: distended abdomen, body condition). In the FRIGHT model we found that gait disorders, kyphosis, and piloerection had the highest contributions to age prediction in our dataset, followed by distended abdomen and body condition (Figure S2B)– all items that our video features were able to predict the score for (Figure 4C). These results together showed that most of the information for aging and frailty came from a small subset of manual FI items and that we are able to predict the information in this subset with video data. Further, as we were able to predict age more accurately and precisely with video data than with manual FI items, video data may also contain additional signals for aging.

Next, we addressed the goal of a vFI (Figure 4A, Model vFI): prediction of manual FI score with video data. Similarly to the vFRIGHT modeling, the random forest regression model predicted FI score on unseen future data better than all other models, with a lowest mean absolute error (MAE) (*p <* 2.1*e*^−15^,F_3,147_ = 30.53), root-mean-squared error (RMSE) (*p <* 8.3*e*^−14^, F_3,147_ = 26.62), and highest R^2^ (*p <* 4.7*e*^−14^, F_3,147_ = 27.2) (Figure 4D, Figure S2E). The model could predict the FI score within 0.04±0.002 of the actual FI score (FI scores have a possible range of 0 to 1, in our dataset we find a range of 0.04 to 0.47). This error is akin to 1 FI item mis-scored at 1 or 2 items mis-scored at 0.5 and demonstrates the robustness of the model. We concluded that our video-generated features can be successfully used for automated frailty scoring. Age has a correlation with manual FI score of r=0.81 which is higher than any video feature. Thus, when we use a model with only age as feature, we find a higher prediction accuracy (Figure S5A). The model using both video features and age (All_*RF*_) does notably better than the model with age alone, showing that the video features provide important information about frailty (Figure S5A). When we looked specifically at mice whose FI scores deviated from their age group– younger mice with higher frailty and older mice with lower frailty, the vFI model (Video_*RF*_) performs noticeably better than the models using age and even the model using video features + age (All_*RF*_) (Figure S5B). This shows that for mice who are outliers of their age group, video features provides better information than age.

Finally, to see how much training data is realistically needed for high performance prediction with vFI and vFRIGHT, we performed a simulation study where we allocated different percentage of total data to training (Figure S5E). We found that a training set of <80% of our current dataset achieved similar performance while a decrease in the training set size below this shows a general downward trend in performance. As open field tests are sometimes done with lengths shorter than 1 hour, we next investigated the decrease in accuracy for vFI predictions using videos with shorter lengths by truncating videos to the first 5 and first 20 minutes (Figure S5D). The features associated with 60-minute videos had the best accuracy for vFI prediction (LMM where ‘simulation’ is the random effect; lowest MAE, F_2,98_ = 178.39, *p <* 2.2*e*^−16^; lowest RMSE, F_2,98_ = 156.93, *p <* 2.2*e*^−16^); highest *R*^2^ (*p <* 2.2*e*^−16^, F_2,98_ = 297.3). We observed a significant drop in performance accuracy when the open field test length is reduced from 60 to 20-minute video (LMM with post hoc pairwise comparisons - MAE, *t*_98_ = 14.82, FDR-adjusted *p <* 0.0001; RMSE, *t*_98_ = 13.69, FDR-adjusted *p <* 0.0001; *R*^2^, *t*_98_ = −19.22, FDR-adjusted *p <* 0.0001). Based on our experiments, we concluded that while 60-minute video-generated features provide the most accurate vFI predictions there is not a substantial loss in accuracy in predictions even with 80% of the videos.

### 3.5 Quantification of uncertainty in frailty index predictions

In addition to quantification of an average accuracy, we investigated prediction errors more closely within our data set in order to see how performance changes across frailty and age. We quantified the prediction error by providing prediction intervals (PIs) that give a range of values, containing the unknown age and FI score with a specified level of confidence, based on the same data that gives random forest point predictions [44]. One existing approach for obtaining random forest-based prediction intervals involves modeling the conditional distribution of FI given the features using quantile random forests [45, 46]. For mice in the test set, we use generalized random forests based on quantiles to provide the point predictions of the FI score (Age resp.) and prediction intervals, which give a range of FI (Age resp.) values that will contain the unknown FI scores (resp. Age) with 95% confidence (Figure S2G,H). The average PI width for all test mouse’s predicted FI score is 0.18±0.04 (resp. 71.96±18.52 for predicted Age), while the PI lengths range from 0.08 to 0.29 (resp. 28 to 113 for Age), highlighting that the widths of the PIs are mouse and age-group specific. We plotted a smoothed regression fit for PIs’ width versus age that indicated the widths increased with the mouse’s age (Figure 4E). The variability of 95% PI widths (Figure 4E) showed higher variability for mice belonging to the middle age groups (labeled M in green). We went beyond simple point predictions by providing prediction intervals (PIs) of the frailty index to quantify our predictions’ uncertainty. This allowed us to pinpoint the FI score and age with higher accuracy for some mice than others.

### 3.6 Feature Importance for Prediction of Frailty

A useful vFI should depend on several features that can capture the mouse’s inherent frailty and simultaneously be interpretable. Interpretability in machine learning is important to prevent bias and instill trust in complex statistical models [47, 48]. In addition, interpretation can guide designs of new features and improve later iterations of the vFI. We took two approaches to identify features important for making vFI predictions using the trained random forest model: (1) feature importance and (2) feature interaction strengths. The feature importance provides a measure of how often the random forest model uses the feature at different depths in the forest. A higher importance value indicates that the feature occurs at the top of the forest and is thus crucial for building the predictive model. For the second approach, we measured a total interaction measure that tells us to what extent a feature interacts with all other model features.

A comparison of the feature importance’s for the vFI and vFRIGHT models (Figure 4A) shows that though many of the most important video features to the model are shared, there are a couple key differences (Figure S5C). For example, step width IQR is much more important for the vFI than for vFRIGHT, and tip-tail lateral displacement (LD) IQR is much more important for vFRIGHT than for vFI. We next obtained a more complete picture of the feature importance by modeling three different quantiles of the FI score’s conditional distribution. The three quantiles represent three frailty groups: low frail (Q1), intermediate frail (M), and high frail (Q3) mice. We hypothesized that different sets of features are crucial for mice belonging to different frailty groups. Indeed, step length1 IQR was crucial in mice belonging to both Q1 (green) and Q3 (red) quantiles (Figure 5A). In addition, features such as length, rear paw speed, dAC/dB (non-gait) and step width were important for lower frailty mice whereas step lengths dB and rear count were more important for animals with high frailty. Similarly, step width, tip tail LD, and width were critical for mice with an FI score close to M (blue).

**Figure 5:**
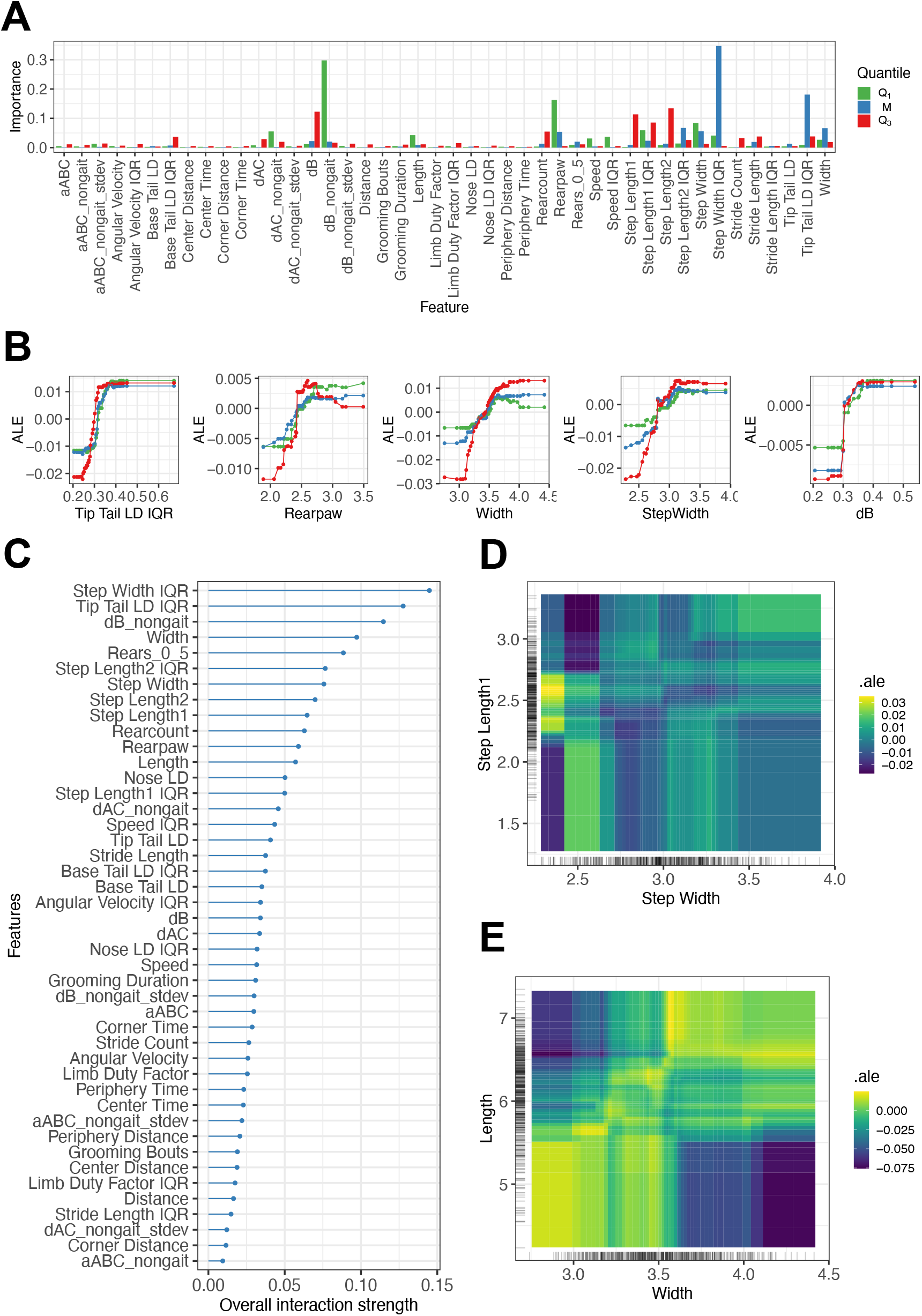
Quantile regression modeling of vFI using Generalized Random Forests. (A) Variable importance measures for three quantile random forest models (lower tail - *Q*_.025_, median - *Q*_.50_,upper tail - *Q*_.975_). Mice in lower and upper tail correspond to mice with low and high frailty scores respectively. (B) Marginal ALE plots show how important features influence the predictions of our models on average. For example, the average predicted FI score rises with increasing step width, but falls for values greater than 3 in mice belonging to lower and upper tail. (C) A plot showing how strongly features interact with each other. (D,E) ALE second-order interaction plots for step width and step length1 (E: Width and Length) on the predicted FI score. Lighter shade indicates an above average and darker shade a below average prediction when the marginal effects from features are already taken into account. Plot D (resp. E) reveals a weak (resp. strong) interaction between step width and step length1 (resp. width and length). Large step width and step length1 increases the vFI score.

For the feature interaction strength approach, we used the H-statistic [49] as an interaction metric that measures the fraction of variability in predictions explained by feature interactions after considering the individual features (Figure 5C). For example, we can explain ≈15% of the prediction function variability due to interaction between step width IQR and other features after considering the individual contributions due to step width IQR and other features. We can explain about 13% and 8% of the prediction function variability due to interaction between tip tail LD IQR and dB non-gait, respectively and other features. We dove deeper and inspected all the two-way interactions between tip tail LD and the other features (results not shown). We found strong tip tail LD interactions with width, stride length, rear paw, and dB of the mouse.

Both feature importance and feature interaction strengths informed us that the trained random forest for vFI depends on several features and their interactions. However, they did not tell us how the vFI depends on these features and how the interactions look. We used the accumulated local effect (ALE) plots [50] that describe how features influence the random forest model’s vFI predictions on average (Figure 5B). For example, an increasing tip tail lateral displacement positively impacts (increases) the predicted FI score for mice in all groups. Similarly, an increasing rear paw measure positively affects the predictions. Mice with larger widths positively affect predictions; larger step widths and dBs positively impact model predictions. Thus the ALE plots for important features provide clear interpretations that are in agreement with our intuition. We explored the ALE second-order interaction effect plot for the step length1-step width (Figure 5D) and Body Length-Width (Figure 5E) predictors. It revealed the two features’ additional interaction effects and did not include the main features’ marginal effects. Figure 5D revealed an interaction between step width and step length: mice with the lowest step widths and step length1 between 2.2 and 2.7 have a higher vFI on average (yellow area) compared to mice with lower step lengths (dark blue area). Similarly, larger widths (3.7 - 4.5) and smaller lengths (4.5 - 5.5) had a negative impact on the average FI scores predictions (Figure 5E).

To summarize, we established vFI’s utility by demonstrating its dependence on several features through marginal feature importance and feature interactions. Next, we used the ALE plots to understand the effects of features on the model predictions, which help us relate the black-box models’ predictions to our video-generated features, an essential final step in our modeling framework.

## 4 Discussion

The mouse FI is an invaluable tool in the study of biological aging. Here, we sought to extend it by producing a scalable automated visual frailty index (vFI) using video-generated features to model FI score. We generated one of the largest frailty data sets for the mouse with associated open field video data. We used machine vision techniques to extract an array of features, many of which show strong correlations with aging and frailty. We also analyzed sex-specific aging in mice. We then trained machine learning models that can accurately predict age and frailty from video features.

We collected our data at a national aging center with similar design as one would in a high-throughput interventional study that may run for several years. Mice were tested by the trained scorer who was available; four different scorers were used to FI test different batches of mice. Further, we had some personnel change between batches. These conditions may provide a more realistic example of inter-lab conditions where discussion and refinement would be difficult. We found that 42% of the variability in our data set could be accounted for by scorer, indicating the presence of a tester effect. This variability and affected some items, such as piloerection, more than others. Although previous studies looking at tester effect found good to high inter-reliability between testers in most cases, FI items showing lower inter-reliability required discussion and refinement for improvement [10, 12].

Top-down videos of mice in the open field were processed by previously trained neural-networks to produce an ellipse-fit, segmentation, and pose estimation of the mouse for each frame. These frame-by-frame measures were used to engineer features. The first category of features were standard open field metrics tracking locomotion, location, and grooming. These metrics had poor correlation with both FI score and age, suggesting that standard open field measures are inadequate to study aging.

In humans, changes in age related body composition and anthropometric measures such as waist-to-hip ratio are predictors of health conditions and mortality risk [51–53]. The effect of aging on body composition in rodent models is less established, though there are observed changes in body composition similar to humans [52, 54]. We found high correlation between morphometric features and both FI score and age, in particular median width and median rear paw width.

Prevalence of gait disorders increase with age [36]. Geriatric patients are shown to have gait irregularities; for example, older adults have increased step width variability [37]. We looked at the spatial, temporal, and postural characteristics of gait for each mouse and found many features with strong correlation to both frailty and age. Analogous to human data, we found a decrease in stride speed with age, as well as an increase in step width variability [37]. As gait is thought to have both cognitive and muscular-skeletal components, it is a compelling area for frailty research.

Spinal mobility in humans is a predictor of quality of life in aged populations [55]. Surprisingly, though some spinal bend metrics showed moderately high correlations with FI score, the relationship was the opposite of what we initially hypothesized. As these metrics are a general account of all the activity of the spine, they are likely capturing a combination of behaviors and body composition which gave this result. Nevertheless, some of these metrics showed a moderately high correlation with FI score and age and were deemed important features in the model.

Many age-related biochemical and physiological changes are known to be sex-specific [30–32]. In humans, there is a known ‘sex-frailty paradox’, where women tend to be more frail but paradoxically live longer [29]. In C57BL/6J mice however, the evidence is mixed [2, 29, 56]. We found that more males survived to old age, leading to a somewhat sex-imbalanced dataset for the oldest mice, and further, we found females tended to have slightly lower frailty distributions than males of the same age group. These results suggest that in C57BL/6J mice, the sex-frailty paradox may not exist or may be reversed. We also found a number of starkly different correlations to age and FI score between males and females. As female mice aged, their tail lateral displacement in strides significantly increased, while males saw almost no change. Males on the other hand showed a strong decrease in stride length with age, while females showed very little change. These differences in gait with age are a new insight and the exact mechanisms driving them are still unknown.

The manual FI evaluates a wider range of body systems than the vFI. However, we believe that the complex behaviors we measure contain implicit information about many body systems. We found that in our isogenic dataset, most information in the manual FI was came from a limited subset of index items. Of the 27 manual FI items scored, 18 items had little to no variation in score in our dataset (almost all mice had the same score i.e. 0), and only 9 items had a balanced distribution of scores. Our video features can accurately predict those 9 FI items. Our model using video features also predicted age more accurately with much less variance than the model using manual FI items (FRIGHT vs. vFRIGHT). This suggest that our video features not only can predict the relevant FI items but also contain signals for aging beyond the traditional manual FI. In addition, the detail in measurements of the features compared to FI items (using actual values rather than a simplified score of 0, 0.5, or 1) could contribute to greater performance.

Finally, using the video features as input to the random forest model, we were able to predict the manual FI score within 0.04±0.002 of the actual score on average, an error akin to 1 FI item mis-scored at 1 or 2 items mis-scored at 0.5. Furthermore, we went beyond simple point predictions by providing 95% prediction intervals. We then applied quantile random forests to low and high quantiles of the FI score’s conditional distribution that revealed how certain features affected frail and healthy animals differently.

Ease of use of the trained model by non-computational labs is an important challenge. Therefore, in addition to implementation details in the Methods section, we have detailed our integrated mouse phenotyping platform– a hardware and software solution that provides tracking, pose estimation, feature generation, and automated behavior analysis in [57]. This platform requires a specific open field apparatus, however, researchers would be able to use the trained model if they generate the same features as our model using their own open field data-collection apparatus. Any set-up that allows tracking and pose estimation using available software would allow researchers to calculate the features necessary to use our trained model.

The vFI can be further improved with the addition of new features through reanalysis of existing data and future technological improvements to data acquisition [58, 59]. For instance, quantification of defecation and urination could provide information about additional systems, while higher camera quality could provide detailed information about fine motor movements based behaviors and appearance based features such as coat condition. Additionally, this approach could potentially be used in a long-term home cage environment. Not only would this further reduce handling and environmental factors, features such as social interaction, feeding, drinking, sleep and others could be integrated. In addition, given the evidence of a strong genetic component to aging [60], application of this method to other strains and genetically heterogeneous populations such as Diversity Outcross and Collaborative Cross, may reveal how genetic variation influences frailty. Further, as predicting mortality risk is a vital function of frailty, video features could be used for studying lifespan. We also imagine that the value of this work could go beyond community adoption and toward community involvement; training data from multiple labs could provide an even more robust and accurate model. This could provide a uniform FI across studies. Overall, our approach has produced novel insights into mouse frailty and shows that video data of mouse behavior can be used to quantify aggregate abstract concepts such as frailty. Our work enables high-throughput, reliable aging studies, particularly interventional studies that are a priority for the aging research community.

## 5 Methods

### 5.1 Mice

C57BL/6J mice were obtained from the Nathan Shock Center at the Jackson Laboratory.

### 5.2 Open Field Assay and Frailty Indexing

The open field behavioral assays were conducted as described in [16, 23, 57]. Mice were shipped from Nathan Shock Center aging colony which resides in different room in the same animal facility at JAX. The aged mice acclimated for 1 week to the Kumar Lab animal holding room, adjacent to the behavioral testing room. During the day of the open field test, mice were allowed to acclimate to the behavior testing room for 30–45 minutes before the start of test. One hour open field testing was performed as previously described [16, 23]. After open field testing mice were returned to the Nathan Shock Center for manual FI. Manual FI was performed by trained experts in the Shock Center within 1 week of the open field assay on each mouse according to previously described protocols[2, 33]. FI testing sheet with all items can be found in Supplementary Materials.

### 5.3 Video, Segmentation, and Tracking

Our open field arena, video apparatus, and tracking and segmentation networks are as detailed previously [23, 57]. Briefly, the open field arena measures 20.5” by 20.5” with Sentech camera mounted 40 inches above. The camera collects data at 30 fps with a 640×480px resolution. We use a neural network trained to produce a segmentation mask of the mouse to produce an ellipse fit of the mouse at each frame as well as a mouse track.

### 5.4 Pose Estimation and Gait

The 12-point 2D pose estimation was produced using a deep convolutional neural network trained as detailed in [25]. The points captured are nose, left ear, right ear, base of neck, left forepaw, right forepaw, mid spine, left rear paw, right rear paw, base of tail, mid tail and tip of tail. Each point at each frame has an x coordinate, a y coordinate, and a confidence score. We use a minimum confidence score of 0.3 to determine which points are included in the analysis.

The gait metrics were produced as detailed in Shephard et al. (2020) [25]. Briefly, the stride cycles were defined by starting and ending with the left hind paw strike, tracked by the pose estimation. These strides were then analyzed for several temporal, spatial, and whole-body coordination characteristics, producing the gait metrics over the entire video.

### 5.5 Open field measures and Feature Engineering

Open field measures were derived from ellipse tracking of mice as described before [23, 24, 57]. The tracking was used to produce locomotor activity and anxiety features. Grooming was classified using a action detection network as previously described [24]. The other engineered features (spinal mobility, body measurements, and rearing) were all derived using the pose estimation data. The spinal mobility metrics used 3 points from the pose: the base of the head (A), the middle of the back (B) and the base of the tail (C). For each frame, the distance between A and C (dAC), the distance between point B and the midpoint of line AC (dB), and the angle formed by the points A,B, and C (aABC) were measured. The means, medians, maximum values, minimum values, and standard deviations of dAC, dB, and aABC were taken over all frames and over frames that were not gait frames (where the animal was not walking). For morphometric measures, we measured the distance between the two rear paw points at each frame and too the means, medians, and standard deviations of that distance over all frames.

For rearing, we took the coordinates of the boundary between the floor and wall of the arean (using OpenCV contour) and added a buffer of 4 pixels. Whenever the mouse’s nose point crossed the buffer, this frame was counted as a rearing frame. Each uninterrupted series of frames where the mouse was rearing (nose crossing the buffer) was counted as a rearing bout. The total number of bouts, the average length of the bouts, the number of bouts in the first 5 minute, and the number of bouts within minutes 5 to 10 were calculated.

### 5.6 Modeling

We investigated the effect of scorer using a linear mixed model with scorer as the random effect and found 42% of the variability (RLRT = 183.85, p *<* 2.2*e*^−16^) in manual FI scores could be accounted for by scorer (Figure 1C). Restricted likelihood-ratio test (RLRT) [61] provided strong evidence of scorer (random) effect with non-zero variance. We fit a cumulative link model (logit link) [62] to the ordinal response (frailty parameter) with weight, age and sex as fixed effects and the tester as a random effect. The effects are estimated variances associated with the random tester effect in the model (Y-axis) across each FI item.

We removed the tester effect from the FI scores using a linear mixed model (LMM) with the lme4 R package [63]. The following model was fit:

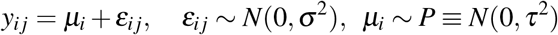

where *y*_*i j*_ is the *j*th animal scored by tester *i, µ*_*i*_ is a tester-specific mean, *ε*_*i j*_ is the animal-specific residual, *σ* ^2^ is the within-tester variance and *P* is the distribution of tester-specific means. We had 4 testers with different number of animals tested by each tester ie *i* = 1, …, 4. The tester effects, estimated with the best linear unbiased predictors (BLUPs) using restricted maximum likelihood estimates [64] were subtracted from the FI scores of the animals, 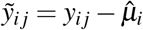.

We modeled tester-adjusted FI scores, 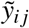, with video-generated features as covariates/inputs using linear regression model with elastic-net penalty [40], support vector machine [41], random forest [42], and gradient boosting machine [43]. We split the data randomly into two parts: train (80%) and test (20%).We ensured that the repeat measurements from the same mouse belonged to either the training or the test data and not both. We used the training data to estimate and tune the models’ hyper-parameters using 10-fold cross-validation; the test set served as an independent evaluation sample for the models’ predictive performance. We performed 50 different splits on the data to allow for a proper assessment of uncertainty in our test set results. The models were compared in terms of mean absolute error (MAE), root-mean-squared-error (RMSE), and *R*^2^. These metrics were compared across the four models using repeated-measures ANOVA through F test with Satterthwaite approximation [65] applied to the test statistic’s denominator degrees of freedom.

For FRIGHT modeling to predict age with manual FI items, we removed frailty parameters with a single value to avoid unstable model fits, i.e., zero-variance predictors. We fit the ordinal regression models [66] without any regularization term and used a global likelihood ratio test (p < 2.2e-16) to determine whether the video features show any evidence of predicting each frailty parameter separately, i.e., evidence of a predictive signal. Next, we used the ordinal regression model with an elastic net penalty [40] to predict frailty parameters using video features.

For predicting manual FI items, we selected frailty parameters for which *p*_*i*_ *<* 0.80, where *i* is the mode of the parameters’ count distribution. For example, Menace reflex is excluded, since *i* = 1 is the mode for Menace reflex’s count distribution with *p*_1_ *>* 0.95.

We obtained the 100(1 − *α*)% out-of-bag prediction intervals I_*α*_ (**X, C**_**n**_), where **X** is the the vector of covariates and *C*_*n*_ is the training set, via quantile random forests [45] with the grf package [46]. Prediction intervals produced with quantile regression forests often perform well in terms of conditional coverage at or above nominal levels i.e. 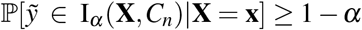 where we set *α* = 0.05.

We picked animals whose ages and FI scores had an inverse relationship, i.e., younger animals with higher FI scores and older animals with lower FI scores. We formed 5 test sets containing animals with these criteria and trained the random forest (RF) model on the remaining mice. We evaluated the predictive accuracy for predicting FI scores for the 5 test sets and displayed the results (Figure S5B). We defined the test sets using age_*L*_, age_*U*_, FI_*L*_, and FI_*U*_, which denote age and FI cutoffs for young and old animals, respectively. For the 5 test sets, we set the parameters as follows,

- age_*L*_ = 60, age_*U*_ = 90, FI_*L*_ = 0.20, and FI_*U*_ = 0.15,
- age_*L*_ = 60, age_*U*_ = 100, FI_*L*_ = 0.20, and FI_*U*_ = 0.15
- age_*L*_ = 50, age_*U*_ = 90, FI_*L*_ = 0.20, and FI_*U*_ = 0.20,
- age_*L*_ = 60, age_*U*_ = 110, FI_*L*_ = 0.20, and FI_*U*_ = 0.20,
- age_*L*_ = 70, age_*U*_ = 100, FI_*L*_ = 0.25, and FI_*U*_ = 0.15

### 5.7 Data and Code Availability

Code and models will be available in Kumar Lab Github account (https://github.com/KumarLabJax and https://www.kumarlab.org/data/). The markdown file in the Github repository https://github.com/KumarLabJax/vFI-modeling contains details for reproducing results in the manuscript and training your own models for vFI/Age prediction. Our manual FI scores and vFI features for all mice in the dataset can be found there as well. Code for engineered features will be found on https://github.com/KumarLabJax/vFI-features.

### 5.8 Supplementary Materials

1. Supplementary Table. Provides breakdown of the number of mice in each batch of testing.
2. Supplementary Notes, merged PDF. Contains 4 tables (detailing video metrics, presenting correlations between vFI features with FI score, presenting correlations between vFI features with age, and presenting correlations between manual FI items with age) and 4 figures.
3. Supplementary Methods, excel sheet. Details the manual FI scoring (items and score explanations)
4. Supplementary Video 1: Young mouse (8 weeks old). Sample of open field video of a mouse aged 8 weeks.
5. Supplementary Video 2: Old mouse (134 weeks old). Sample of open field video of a mouse aged 134 weeks.
6. Supplementary Video 3: Flexibility metrics. At each frame, 3 points shown in yellow are estimated: base of head (A), mid-back (B), and base of tail (C). At each frame, the distance between points A and C (dAC, shown in red), distance between point B and the midpoint of line AC (dB, shown in green), and the angle formed by the points ABC (aABC, shown in blue) are calculated.
7. Supplementary Video 4: Rearing metrics. Rearing is called when the nose of the mice (marked with a red dot) crosses the perimeter of the open field (yellow line). Rearing is indicated by the presence of the red square in the upper corner of video.

## 7 Acknowledgements

We thank Kumar Lab members, Sean Deats, Tom Sproule, Brian Geuther, and Keith Sheppard for behavioral testing, data processing, and helpful advice. We thank Shock Center and Churchill Lab members Hannah Donato, Gaven Garland, Mackenzie Leland, and Laura Robinson for frailty indexing and coordinating. We thank Taneli Helenius for editing. We thank JAX Information Technology team members Edwardo Zaborowski, Shane Sanders, Rich Brey, David McKenzie, and Jason Macklin for infrastructure support. This work was funded by The Jackson Laboratory Directors Innovation Fund, National Institute of Health DA041668 (NIDA, V.K.), DA048634 (NIDA, V.K.). and Nathan Shock Centers of Excellence in the Basic Biology of Aging AG38070(NIA, G.C.).

---

**Table S1:** Testing batches

| Batch | Males | Females | Total Mice | Repeated Mice |
| --- | --- | --- | --- | --- |
| Batch 1 | 141 | 81 | 222 | NA |
| Batch 2 | 173 | 146 | 319 | 105 |
| Batch 3 | 57 | 45 | 102 | 33 |
| Total | 371 | 272 | 643 |  |

## 8 Supplementary Materials

**Table S2:** Video Features

| Video Metrics |  |  |  |
| --- | --- | --- | --- |
| Category | Name | Description | Units |
| Open Field | distance_cm | Sum of locomotor activity. | cm |
| Open Field | center_time_secs | Sum of time spent in center. | sec |
| Open Field | periphery_time_secs | Sum of time spent along any wall. | sec |
| Open Field | corner_time_secs | Sum of time spent in any corner. | sec |
| Open Field | center_distance_cm | Average distance from center across the video. | cm |
| Open Field | periphery_distance_cm | Average distance from nearest periphery across the video. | cm |
| Open Field | corner_distance_cm | Average distance from nearest corners across the video. | cm |
| Open Field | grooming_number_bouts | Sum of all grooming bouts in video. | ~ |
| Open Field | grooming_duration_secs | Average length of grooming bouts. | sec |
| Gait | angular_velocity | The first derivative of angle of a mouse, determined by the vector connecting the mouse's base of tail to its base of neck | deg/second |
| Gait | lateral_displacement | The difference between the minimum and maximum values of a reference point's perpendicular distance from the mouse's displacement vector for a stride for each frame of a stride, normalized by the mouse's body length. The reference points used are nose, base of tail, and tail tip. | ~ |
| Gait | limb_duty_factor | The amount of time that the paw is in contact with the ground divided by the full stride time, calculated and averaged for each hind paw. | ~ |
| Gait | speed_cm_per_sec | Speed is determined by the base of tail point. | cm/second |
| Gait | step_length | The distance that the right hind paw travels past the previous opposite paw strike. Step_length 1 uses left hind paw strike, while step_length2 uses right hind paw strike. | cm |

| Video Features |  |  |  |
| --- | --- | --- | --- |
| Category | Name | Description | Units |
| Gait | step_width | The length of the shortest line segment that connects the right hind paw strike to the line that connects the left hind paw's toe-off location to its subsequent foot strike position. | cm |
| Gait | stride_length | The full distance that the left hind paw travels for a stride, from toe-off to foot-strike. | cm |
| Gait | temporal_symmetry | The difference in time between the left and right hindpaw strike, divided by the total strike time. | ~ |
| Gait | stride_count | Sum of all recorded strides in video. | strides |
| Gait | distance_cm_sc | Sum of locomotor activity, normalized by time spent in open field. | cm/second |
| Engineered | dAC | Distance between base of head and base of tail, normalized by the max dAC recorded. | cm |
| Engineered | dB | The distance between the mid-back point and the midpoint of the line AC. | cm |
| Engineered | aABC | The angle between the base of head point, mid-back point, and base of tail point. | degrees |
| Engineered | width | Width of the ellipse fit for the mouse calculated for all frames. | cm |
| Engineered | length | Length of the ellipse fit for the mouse calculated for all frames. | cm |
| Engineered | rearpaw | The distance between rearpaws calculated for all frames. | cm |
| Engineered | rear_count | Sum of rearing bouts in video. | ~ |
| Engineered | avg_rear_len | Average length of rearing bouts in video. | sec |
| End of Table |  |  |  |

**Table S3:** Feature Correlation with FI score

| vFI feature correlation (Pearson) with FI score |  |  |  |  |  |  |
| --- | --- | --- | --- | --- | --- | --- |
| Features | All Mice |  | Males |  | Females |  |
|  | Cor | Pvalue | Cor | Pvalue | Cor | Pvalue |
| tip_tail_lateral_displacement_iqr | 0.629 | 0.000 | 0.582 | 0.000 | 0.659 | 0.000 |
| median_step_width | 0.561 | 0.000 | 0.513 | 0.000 | 0.559 | 0.000 |
| median_width | 0.556 | 0.000 | 0.561 | 0.000 | 0.497 | 0.000 |
| step_length2_iqr | 0.551 | 0.000 | 0.595 | 0.000 | 0.421 | 0.000 |
| base_tail_lateral_displacement_iqr | 0.547 | 0.000 | 0.498 | 0.000 | 0.614 | 0.000 |
| median_rearpaw | 0.545 | 0.000 | 0.493 | 0.000 | 0.552 | 0.000 |
| step_length1_iqr | 0.544 | 0.000 | 0.584 | 0.000 | 0.418 | 0.000 |
| step_width_iqr | 0.504 | 0.000 | 0.474 | 0.000 | 0.507 | 0.000 |
| median_length | 0.494 | 0.000 | 0.386 | 0.000 | 0.552 | 0.000 |
| stride_length_iqr | 0.481 | 0.000 | 0.483 | 0.000 | 0.425 | 0.000 |
| avg_bout_len | -0.467 | 0.000 | -0.456 | 0.000 | -0.448 | 0.000 |
| temporal_symmetry_iqr | 0.458 | 0.000 | 0.485 | 0.000 | 0.350 | 0.000 |
| dB_median | 0.455 | 0.000 | 0.382 | 0.000 | 0.496 | 0.000 |
| median_speed_cm_per_sec | -0.442 | 0.000 | -0.493 | 0.000 | -0.335 | 0.000 |
| dB_nongait_median | 0.429 | 0.000 | 0.328 | 0.000 | 0.561 | 0.000 |
| dAC_nongait_median | 0.402 | 0.000 | 0.320 | 0.000 | 0.440 | 0.000 |
| median_stride_length | -0.395 | 0.000 | -0.480 | 0.000 | -0.189 | 0.002 |
| rears_0_5 | -0.390 | 0.000 | -0.362 | 0.000 | -0.391 | 0.000 |
| dAC_stdev | -0.317 | 0.000 | -0.341 | 0.000 | -0.225 | 0.000 |
| limb_duty_factor_iqr | 0.314 | 0.000 | 0.346 | 0.000 | 0.184 | 0.002 |
| dAC_min | 0.290 | 0.000 | 0.323 | 0.000 | 0.232 | 0.000 |
| median_tip_tail_lateral_displacement | 0.275 | 0.000 | 0.126 | 0.012 | 0.558 | 0.000 |
| rear_count | -0.274 | 0.000 | -0.348 | 0.000 | -0.155 | 0.010 |
| speed_cm_per_sec_iqr | -0.247 | 0.000 | -0.325 | 0.000 | -0.165 | 0.006 |
| rears_5_10 | -0.237 | 0.000 | -0.259 | 0.000 | -0.169 | 0.005 |
| aABC_nongait_stdev | -0.227 | 0.000 | -0.361 | 0.000 | -0.063 | 0.297 |
| corner_distance_cm | -0.224 | 0.000 | -0.263 | 0.000 | -0.118 | 0.049 |
| aABC_stdev | -0.222 | 0.000 | -0.340 | 0.000 | -0.097 | 0.108 |
| aABC_min | 0.221 | 0.000 | 0.257 | 0.000 | 0.176 | 0.003 |
| median_step_length1 | -0.216 | 0.000 | -0.293 | 0.000 | -0.014 | 0.823 |
| angular_velocity_iqr | 0.209 | 0.000 | 0.175 | 0.000 | 0.348 | 0.000 |
| grooming_number_bouts | -0.190 | 0.000 | -0.217 | 0.000 | -0.094 | 0.119 |
| dB_nongait_min | 0.186 | 0.000 | 0.191 | 0.000 | 0.214 | 0.000 |
| periphery_distance_cm | -0.178 | 0.000 | -0.246 | 0.000 | -0.039 | 0.521 |
| median_step_length2 | -0.178 | 0.000 | -0.256 | 0.000 | -0.016 | 0.794 |
| distance_cm | -0.161 | 0.000 | -0.229 | 0.000 | -0.027 | 0.655 |
| median_temporal_symmetry | 0.160 | 0.000 | 0.174 | 0.001 | 0.138 | 0.024 |
| grooming_duration_secs | -0.151 | 0.000 | -0.111 | 0.029 | -0.185 | 0.002 |
| median_base_tail_lateral_displacement | 0.149 | 0.000 | -0.003 | 0.950 | 0.511 | 0.000 |
| aABC_median | -0.148 | 0.000 | -0.093 | 0.065 | -0.149 | 0.013 |
| periphery_time_secs | -0.115 | 0.003 | -0.093 | 0.067 | -0.140 | 0.020 |
| Distance cm/sc | -0.105 | 0.006 | -0.182 | 0.000 | 0.068 | 0.259 |
| dB_nongait_max | -0.102 | 0.009 | -0.095 | 0.061 | -0.121 | 0.046 |
| dAC_nongait_stdev | -0.096 | 0.014 | -0.144 | 0.005 | -0.042 | 0.490 |
| nose_lateral_displacement_iqr | -0.095 | 0.013 | -0.164 | 0.001 | 0.121 | 0.045 |

| vFI Feature correlation (Pearson) with FI score (Table continued) |  |  |  |  |  |  |
| --- | --- | --- | --- | --- | --- | --- |
| Features | All Mice |  | Males |  | Females |  |
|  | Cor | Pvalue | Cor | Pvalue | Cor | Pvalue |
| corner_time_secs | -0.087 | 0.024 | -0.013 | 0.803 | -0.164 | 0.006 |
| center_distance_cm | -0.074 | 0.057 | -0.119 | 0.018 | 0.006 | 0.921 |
| dAC_median | 0.071 | 0.066 | 0.047 | 0.352 | 0.131 | 0.030 |
| dB_max | 0.064 | 0.100 | 0.030 | 0.557 | 0.063 | 0.297 |
| aABC_nongait_median | -0.059 | 0.128 | -0.007 | 0.888 | -0.120 | 0.046 |
| center_time_secs | 0.043 | 0.266 | 0.046 | 0.364 | 0.011 | 0.855 |
| dB_stdev | 0.041 | 0.285 | 0.006 | 0.910 | 0.057 | 0.343 |
| stride_count | -0.034 | 0.378 | -0.023 | 0.649 | 0.052 | 0.392 |
| dB_nongait_stdev | -0.031 | 0.432 | -0.080 | 0.117 | 0.030 | 0.615 |
| median_angular_velocity | 0.025 | 0.514 | -0.012 | 0.807 | 0.108 | 0.073 |
| median_limb_duty_factor | 0.007 | 0.853 | -0.033 | 0.517 | 0.092 | 0.127 |
| median_nose_lateral_displacement | -0.006 | 0.885 | -0.034 | 0.501 | 0.160 | 0.008 |
| End of Table |  |  |  |  |  |  |

**Table S4:** vFI feature correlation (Pearson) with age

| Feature Correlation with Age |  |  |  |  |  |  |
| --- | --- | --- | --- | --- | --- | --- |
| Features | All Mice |  | Males |  | Females |  |
|  | Cor | Pvalue | Cor | Pvalue | Cor | Pvalue |
| tip_tail_lateral_displacement_iqr | 0.760 | 0.000 | 0.728 | 0.000 | 0.795 | 0.000 |
| median_step_width | 0.662 | 0.000 | 0.609 | 0.000 | 0.695 | 0.000 |
| median_width | 0.648 | 0.000 | 0.646 | 0.000 | 0.628 | 0.000 |
| step_length2_iqr | 0.638 | 0.000 | 0.686 | 0.000 | 0.503 | 0.000 |
| step_length1_iqr | 0.637 | 0.000 | 0.691 | 0.000 | 0.494 | 0.000 |
| base_tail_lateral_displacement_iqr | 0.637 | 0.000 | 0.595 | 0.000 | 0.707 | 0.000 |
| median_rearpaw | 0.636 | 0.000 | 0.579 | 0.000 | 0.668 | 0.000 |
| step_width_iqr | 0.609 | 0.000 | 0.583 | 0.000 | 0.611 | 0.000 |
| stride_length_iqr | 0.564 | 0.000 | 0.575 | 0.000 | 0.502 | 0.000 |
| avg_bout_len | -0.559 | 0.000 | -0.559 | 0.000 | -0.530 | 0.000 |
| median_length | 0.540 | 0.000 | 0.447 | 0.000 | 0.605 | 0.000 |
| temporal_symmetry_iqr | 0.537 | 0.000 | 0.550 | 0.000 | 0.450 | 0.000 |
| dB_nongait_median | 0.500 | 0.000 | 0.418 | 0.000 | 0.620 | 0.000 |
| dB_median | 0.491 | 0.000 | 0.402 | 0.000 | 0.570 | 0.000 |
| median_speed_cm_per_sec | -0.482 | 0.000 | -0.508 | 0.000 | -0.401 | 0.000 |
| dAC_nongait_median | 0.421 | 0.000 | 0.339 | 0.000 | 0.470 | 0.000 |
| rears_0_5 | -0.420 | 0.000 | -0.380 | 0.000 | -0.442 | 0.000 |
| median_stride_length | -0.411 | 0.000 | -0.499 | 0.000 | -0.170 | 0.005 |
| dAC_stdev | -0.399 | 0.000 | -0.420 | 0.000 | -0.302 | 0.000 |
| dAC_min | 0.381 | 0.000 | 0.390 | 0.000 | 0.359 | 0.000 |
| median_tip_tail_lateral_displacement | 0.351 | 0.000 | 0.191 | 0.000 | 0.698 | 0.000 |
| limb_duty_factor_iqr | 0.334 | 0.000 | 0.379 | 0.000 | 0.171 | 0.004 |
| aABC_min | 0.303 | 0.000 | 0.328 | 0.000 | 0.271 | 0.000 |
| aABC_nongait_stdev | -0.295 | 0.000 | -0.369 | 0.000 | -0.204 | 0.001 |
| aABC_stdev | -0.291 | 0.000 | -0.357 | 0.000 | -0.231 | 0.000 |
| rear_count | -0.274 | 0.000 | -0.313 | 0.000 | -0.192 | 0.001 |
| speed_cm_per_sec_iqr | -0.272 | 0.000 | -0.319 | 0.000 | -0.218 | 0.000 |
| rears_5_10 | -0.260 | 0.000 | -0.248 | 0.000 | -0.240 | 0.000 |
| grooming_number_bouts | -0.244 | 0.000 | -0.241 | 0.000 | -0.184 | 0.002 |
| median_base_tail_lateral_displacement | 0.226 | 0.000 | 0.078 | 0.120 | 0.647 | 0.000 |
| median_step_length1 | -0.215 | 0.000 | -0.288 | 0.000 | 0.010 | 0.868 |
| grooming_duration_secs | -0.209 | 0.000 | -0.172 | 0.001 | -0.247 | 0.000 |
| median_temporal_symmetry | 0.207 | 0.000 | 0.228 | 0.000 | 0.174 | 0.004 |
| angular_velocity_iqr | 0.207 | 0.000 | 0.175 | 0.000 | 0.352 | 0.000 |
| corner_distance_cm | -0.199 | 0.000 | -0.170 | 0.001 | -0.175 | 0.004 |
| dB_nongait_min | 0.179 | 0.000 | 0.231 | 0.000 | 0.149 | 0.013 |
| nose_lateral_displacement_iqr | -0.161 | 0.000 | -0.215 | 0.000 | 0.052 | 0.393 |
| distance_cm | -0.155 | 0.000 | -0.184 | 0.000 | -0.062 | 0.305 |
| dAC_nongait_stdev | -0.152 | 0.000 | -0.200 | 0.000 | -0.093 | 0.125 |
| periphery_distance_cm | -0.152 | 0.000 | -0.168 | 0.001 | -0.071 | 0.239 |
| median_step_length2 | -0.135 | 0.000 | -0.212 | 0.000 | 0.055 | 0.359 |
| aABC_median | -0.130 | 0.001 | -0.090 | 0.074 | -0.108 | 0.074 |
| center_distance_cm | -0.114 | 0.003 | -0.159 | 0.002 | -0.025 | 0.683 |
| Distance cm/sc | -0.092 | 0.017 | -0.147 | 0.003 | 0.071 | 0.237 |
| dB_stdev | 0.077 | 0.048 | 0.069 | 0.171 | 0.031 | 0.611 |

| vFI feature correlation (Pearson) with age (Table continued) |  |  |  |  |  |  |
| --- | --- | --- | --- | --- | --- | --- |
| Features | All Mice |  | Males |  | Females |  |
|  | Cor | Pvalue | Cor | Pvalue | Cor | Pvalue |
| dB_nongait_max | -0.076 | 0.050 | -0.083 | 0.103 | -0.077 | 0.205 |
| corner_time_secs | -0.075 | 0.052 | 0.021 | 0.685 | -0.196 | 0.001 |
| periphery_time_secs | -0.074 | 0.056 | -0.018 | 0.716 | -0.148 | 0.014 |
| dB_nongait_stdev | -0.071 | 0.067 | -0.123 | 0.015 | 0.002 | 0.979 |
| dB_max | 0.070 | 0.070 | 0.058 | 0.247 | 0.015 | 0.805 |
| dAC_median | 0.062 | 0.108 | 0.017 | 0.736 | 0.170 | 0.005 |
| median_nose_lateral_displacement | -0.046 | 0.233 | -0.077 | 0.126 | 0.139 | 0.021 |
| median_angular_velocity | 0.037 | 0.341 | 0.008 | 0.872 | 0.111 | 0.065 |
| aABC_nongait_median | -0.034 | 0.378 | -0.003 | 0.949 | -0.074 | 0.223 |
| stride_count | -0.024 | 0.537 | -0.013 | 0.800 | 0.072 | 0.231 |
| median_limb_duty_factor | -0.022 | 0.571 | -0.042 | 0.406 | 0.041 | 0.500 |
| center_time_secs | -0.002 | 0.958 | -0.033 | 0.512 | 0.012 | 0.846 |
| End of Table |  |  |  |  |  |  |

**Table S5:** Manual FI item correlation with age

| Features | All Mice |  | Males |  | Females |  |
| --- | --- | --- | --- | --- | --- | --- |
|  | Correlation | p-value | Correlation | p-value | Correlation | p-value |
| Piloerection | 0.721 | 0.000 | 0.641 | 0.000 | 0.797 | 0.000 |
| Kyphosis | 0.703 | 0.000 | 0.660 | 0.000 | 0.725 | 0.000 |
| Gait.disorders | 0.674 | 0.000 | 0.631 | 0.000 | 0.694 | 0.000 |
| Coat.condition | 0.661 | 0.000 | 0.675 | 0.000 | 0.597 | 0.000 |
| Body.condition | 0.551 | 0.000 | 0.519 | 0.000 | 0.569 | 0.000 |
| Distended.abdomen | 0.548 | 0.000 | 0.523 | 0.000 | 0.527 | 0.000 |
| Loss.of.fur.colour | 0.444 | 0.000 | 0.442 | 0.000 | 0.326 | 0.000 |
| Tumours | 0.319 | 0.000 | 0.327 | 0.000 | 0.239 | 0.000 |
| Vestibular.disturbance | 0.317 | 0.000 | 0.297 | 0.000 | 0.310 | 0.000 |
| Breathing.rate.depth | 0.307 | 0.000 | 0.290 | 0.000 | 0.319 | 0.000 |
| Righting.Reflex | 0.287 | 0.000 | 0.254 | 0.000 | 0.347 | 0.000 |
| Tremor | 0.280 | 0.000 | 0.212 | 0.000 | 0.381 | 0.000 |
| Tail.stiffening | 0.261 | 0.000 | 0.262 | 0.000 | 0.230 | 0.000 |
| Loss.of.whiskers | 0.244 | 0.000 | 0.276 | 0.000 | 0.370 | 0.000 |
| Vision.loss..Visual.Placing. | 0.213 | 0.000 | 0.189 | 0.000 | 0.334 | 0.000 |
| Alopecia | 0.195 | 0.000 | 0.149 | 0.003 | 0.438 | 0.000 |
| Eye.discharge.swelling | 0.069 | 0.072 | 0.049 | 0.332 | 0.103 | 0.081 |
| Menace.reflex | -0.055 | 0.156 | -0.027 | 0.600 | -0.078 | 0.188 |
| Corneal.opacity | 0.032 | 0.412 | 0.020 | 0.696 | 0.071 | 0.231 |
| Cataracts | 0.007 | 0.860 | -0.020 | 0.692 | 0.055 | 0.351 |
| Dermatitis | 0.007 | 0.862 | 0.042 | 0.410 | -0.009 | 0.875 |
| Malocclusions | -0.007 | 0.863 | 0.009 | 0.860 | -0.038 | 0.521 |
| Microphthalmia | 0.002 | 0.966 | 0.015 | 0.764 | -0.019 | 0.745 |
| Nasal.discharge |  |  |  |  |  |  |
| Rectal.prolapse |  |  |  |  |  |  |
| Vaginal.uterine. |  |  |  |  |  |  |
| Diarrhea |  |  |  |  |  |  |

**Figure S1:**
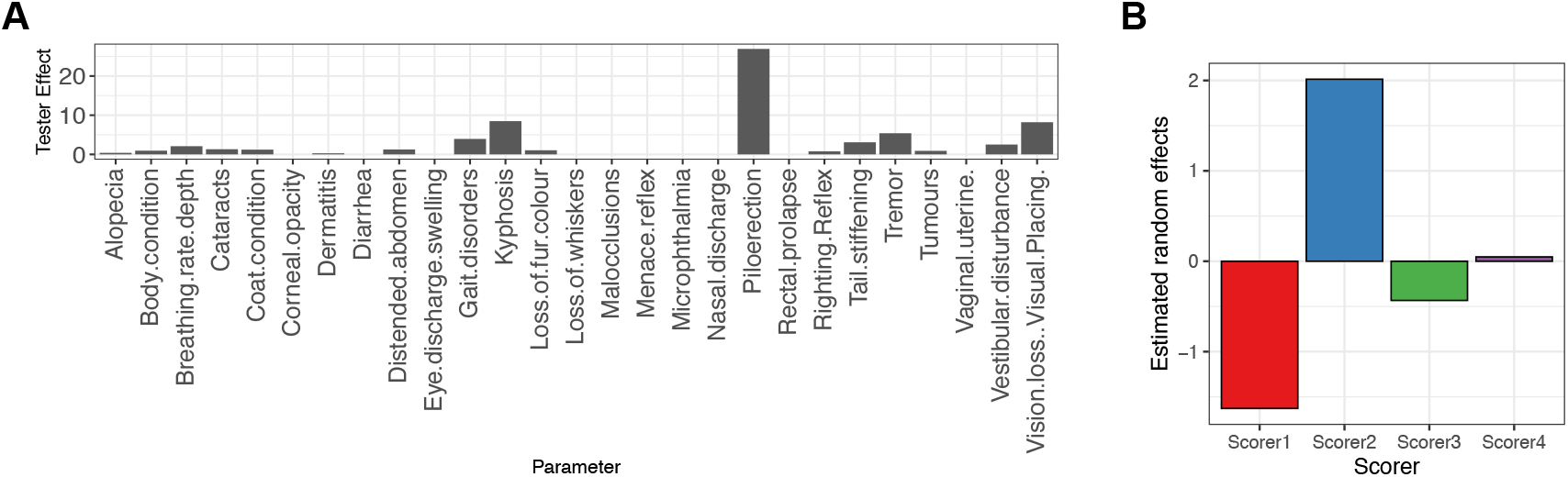
Estimation of the scorer effect in clinical FI items. (A) The effect of tester varies across FI items. (B) The estimated random effect across 4 scorers in the data set.

**Figure S2:**
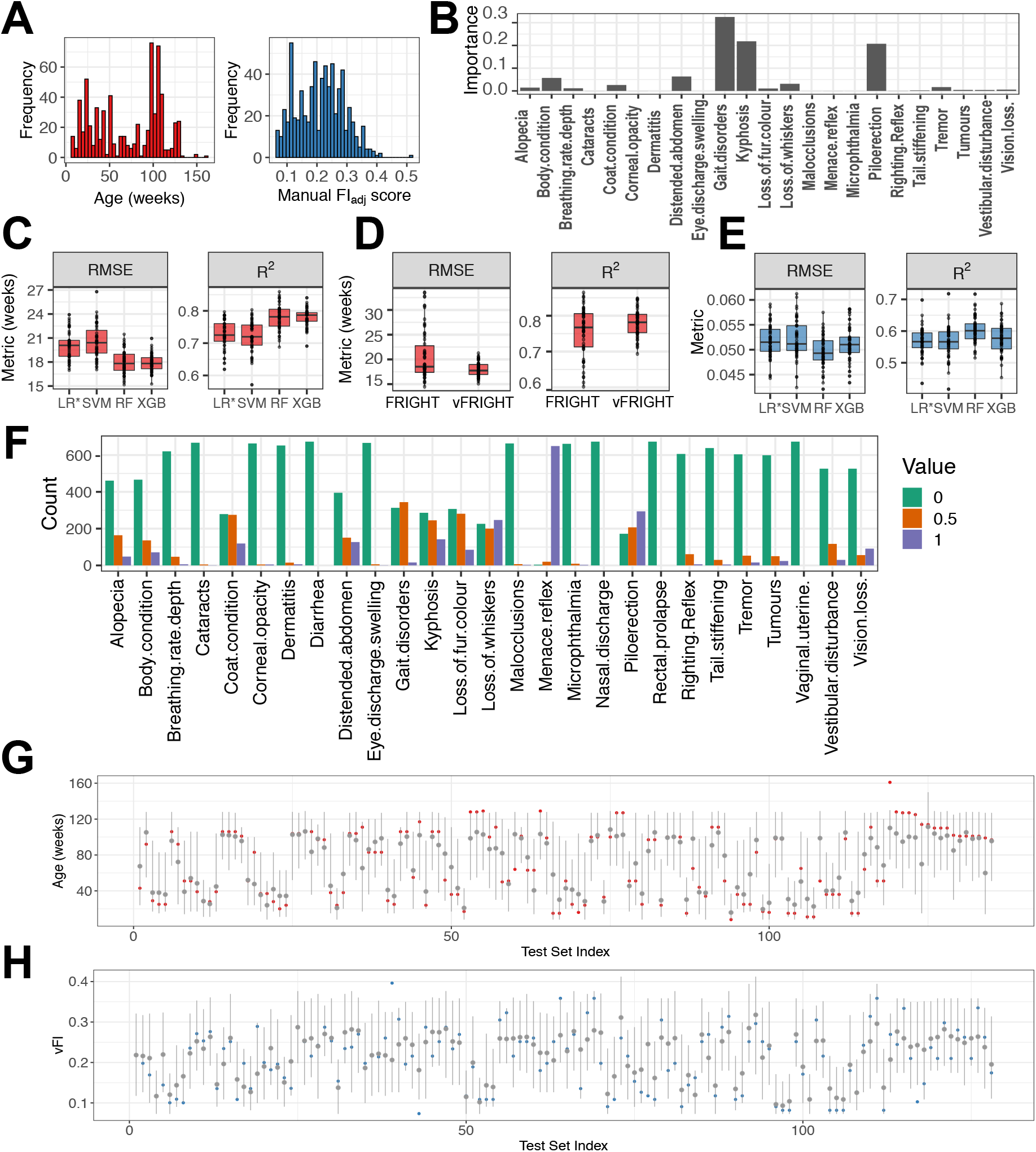
Detailed modeling analysis (A) The distribution of age across 533 mice. The distribution of manual FI_adj_ scores (range 0-0.5) across 533 mice. (B) To determine the contributions of frailty parameters in predicting Age, we calculated the feature importance of all frailty parameters. We discover that gait disorders, kyphosis, and piloerection have the highest contributions. (C) The random forest regression model performed better than other models with the lowest root-mean-squared error (RMSE) (*p <* 2.2*e*^−16^, F_3,147_ = 59.53), and highest R^2^ (*p <* 2.2*e*^−16^, F_3,147_ = 58.14) when compared using repeated-measures ANOVA (D) The vFRIGHT model performed better than the FRIGHT model with a lower RMSE (RMSE_vFRIGHT_ = 17.97 ± 1.44, RMSE_FRIGHT_ = 20.62 ± 4.78, *p <* 6.1*e* − 7, *F*_1,49_ = 32.84) and higher R^2^ (RMSE_vFRIGHT_ = 0.78±0.04, RMSE_FRIGHT_ = 0.76±0.07, *p <* 2.1*e*−8, *F*_1,49_ = 44.54) when compared using repeated-measures ANOVA (E) The random forest regression model for predicting FI score on unseen future data performed better than all other models, with a lowest root-mean-squared error (RMSE) (*p <* 8.3*e*^−14^, F_3,147_ = 26.62), and highest R^2^ (*p <* 4.7*e*^−14^, F_3,147_ = 27.2) (F)The plot shows the counts distribution (0 - green, 0.5 - orange, 1 - purple) for individual frailty parameters— for many parameters such as Nasal discharge, Rectal prolapse, Vaginal uterine, and Diarrhea, the proportion of 0 counts is 1 (*p*_0_ = 1). Similarly, Dermatitis, Cataracts, Eye discharge swelling, Microphthalmia, Corneal opacity, Tail stiffening, and Malocclusions have *p*_0_ *>* 0.95. (G,H) Out-of-bag (OOB) error based 95% prediction intervals (PIs) (gray lines) quantifying uncertainty in point estimates/predictions (gray dots). There is one interval per test mouse, and approximately 95% of the PI intervals contain the correct Age (red dots) and FI scores (blue dots).

**Figure S3:**
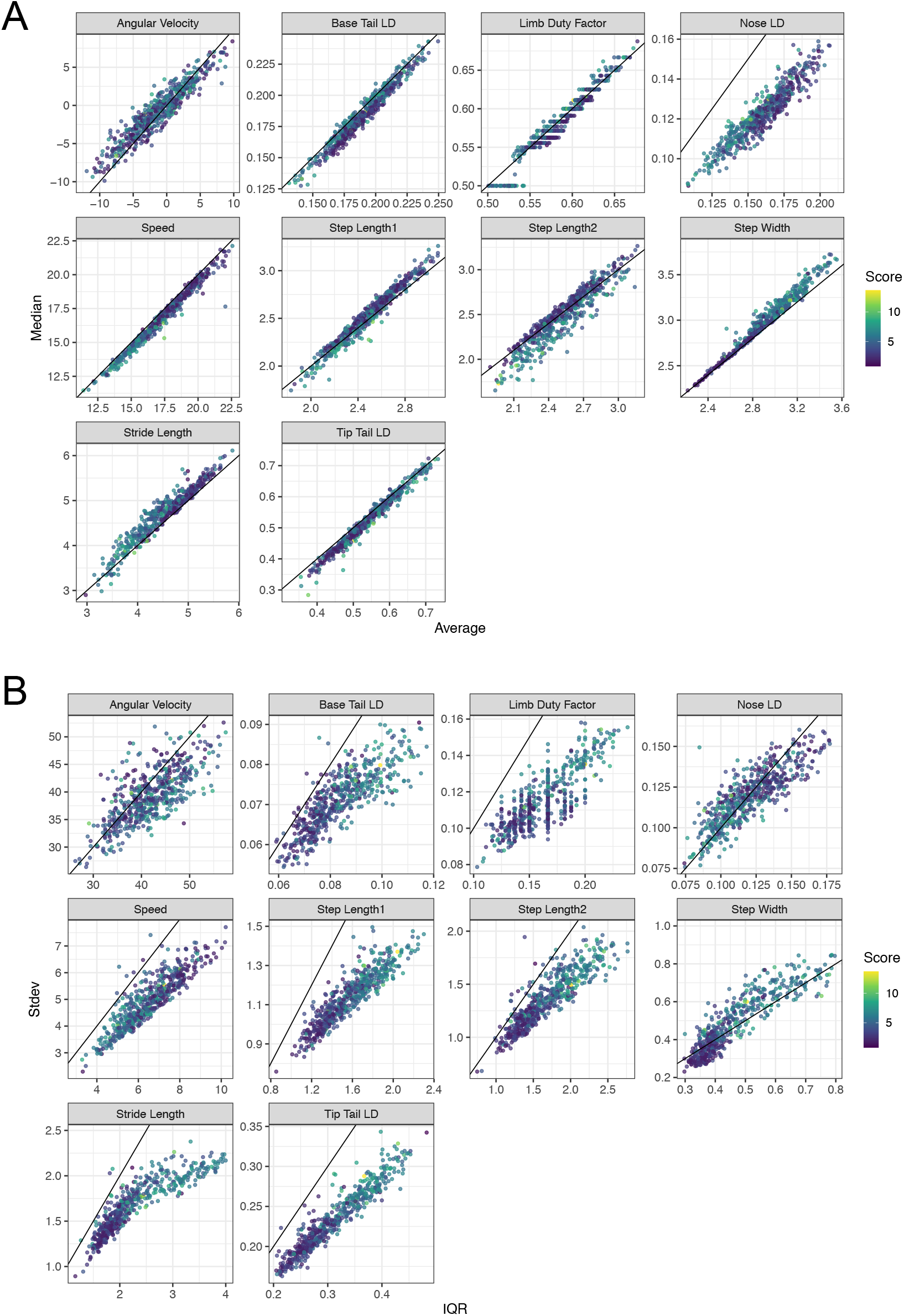
Correlation between video metrics. (A) Correlation between average/mean (x-axis) and median (y-axis) video gait metrics. The diagonal line corresponds to maximum correlation i.e 1. (B) Correlation between inter-quartile range (IQR, x-axis) and standard deviation (Stdev, y-axis) video gait metrics. The diagonal line corresponds to maximum correlation i.e 1. A tight wrap of points around the diagonal line indicates a high correlation between mean and median or IQR and standard deviation for the respective metric.

**Figure S4:**
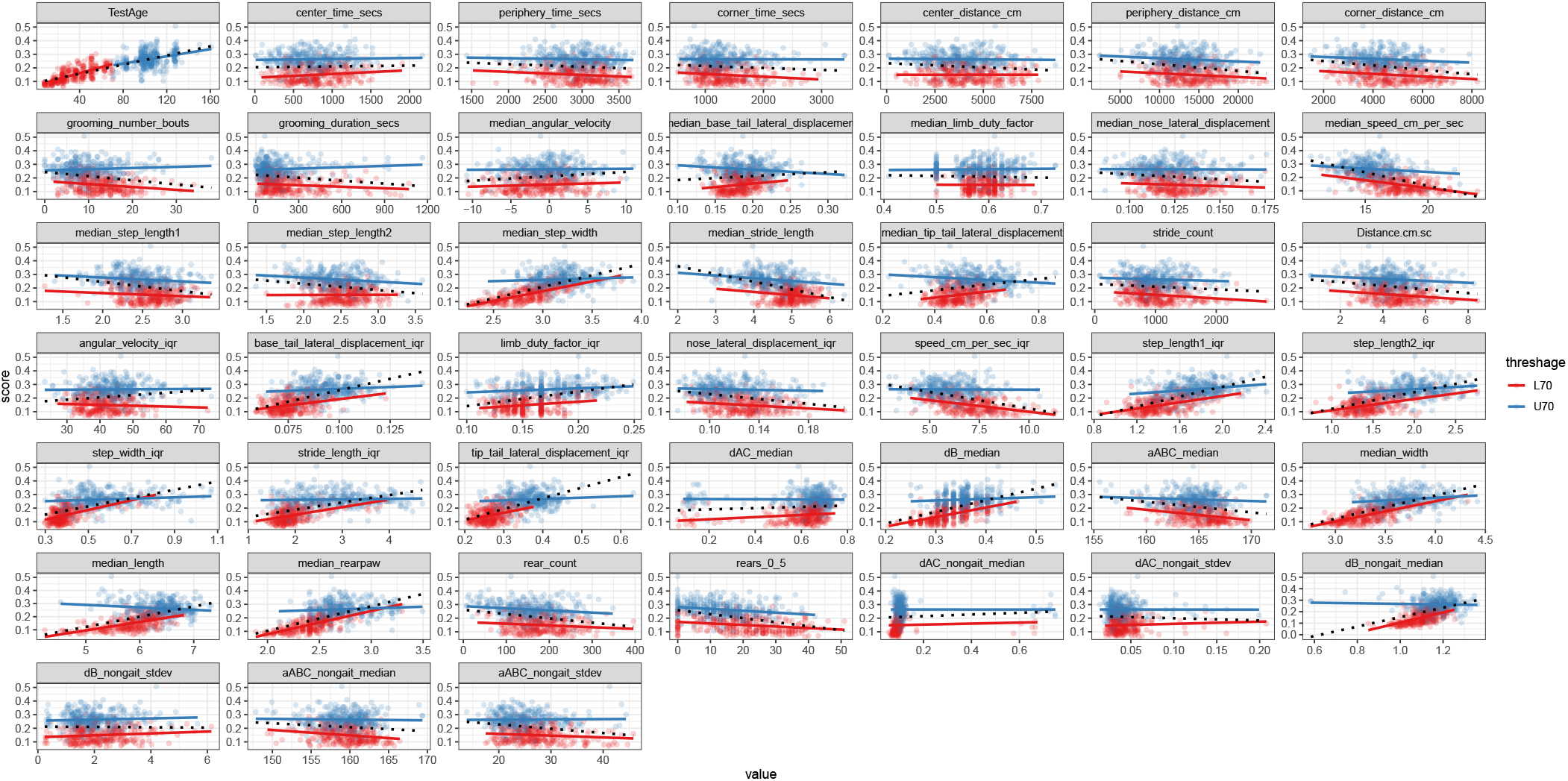
Test for Simpson’s paradox. Simpson [34] showed that the statistical relationship observed in the population could be reversed within all of the subgroups that make up that population, leading to erroneous conclusions drawn from the population data. To test for the manifestation of Simpson’s paradox in our data, we split the bimodal Age distribution into two separate unimodal distributions (clusters), i.e., less than 70 weeks old (L70, red) versus more than 70 weeks old (U70, blue). Next, we plotted the dependent variable (frailty) against each of the independent variables/features in our data and fit a simple linear regression model to each subgroup separately (solid red and blue lines) as well as to the aggregate data (black dotted line). We find no evidence of Simpson’s paradox in our dataset.

**Figure S5:**
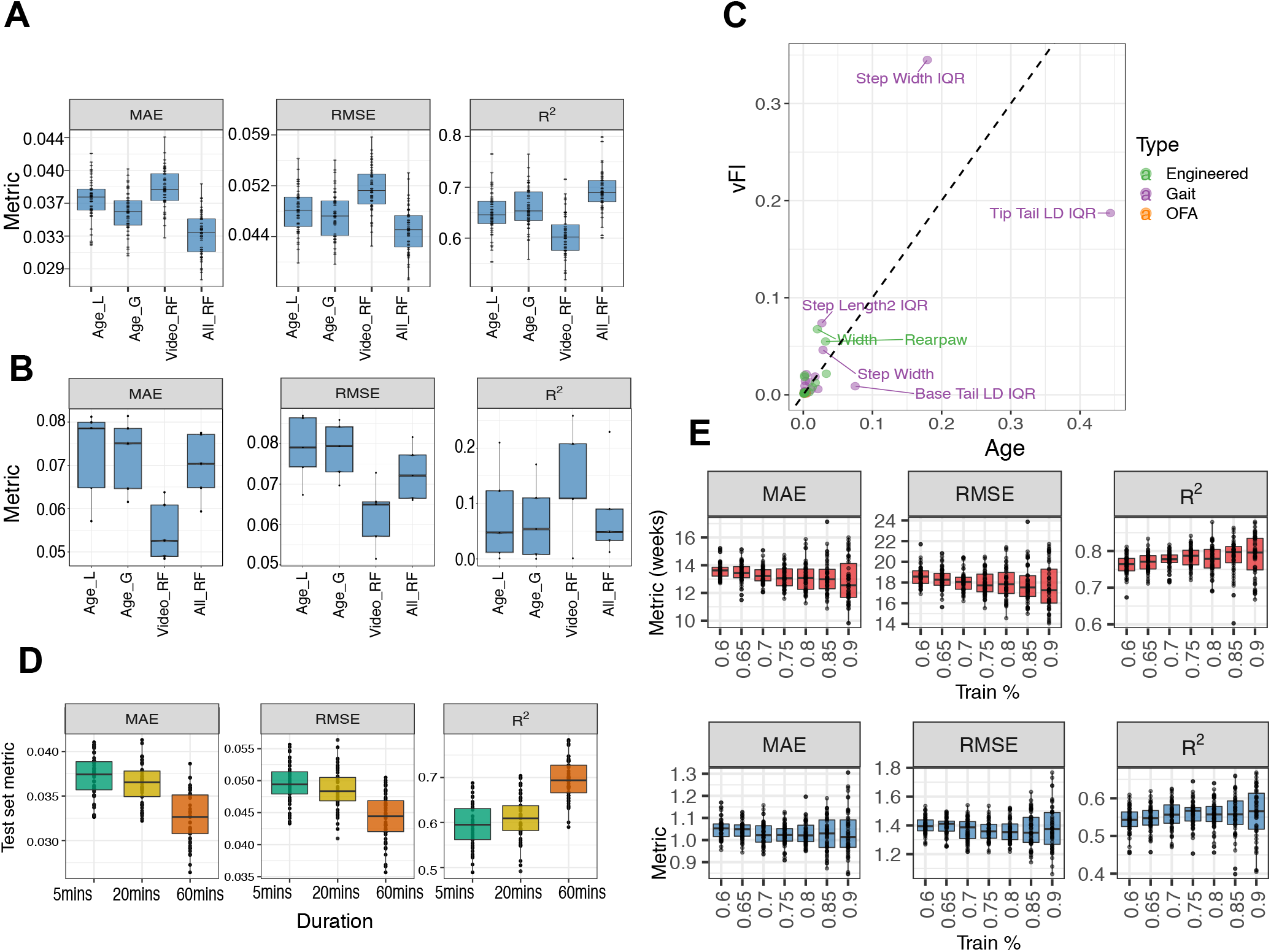
Further experiments to test model performance and parameters. (A) We compare the performance of different feature sets, 1) age alone, 2) video, and 3) age + video, in predicting frailty. We use age alone as a feature in a linear (Age_*L*_) and a generalized additive non-linear model (Age_*G*_). Although we didn’t notice a clear improvement of the random forest model (Video_*RF*_) using video features over a vFI prediction based on age alone, a clear improvement in prediction performance is seen for the model (All_*RF*_), which contains video features + age. This shows that video features add important information pertaining to frailty that age alone does not. (B) We picked animals whose ages and FI scores had an inverse relationship, i.e., younger animals with higher FI scores and older animals with lower FI scores. We formed 5 test sets containing animals with these criteria and trained the random forest (RF) model on the remaining mice. The model using only video features (Video_*RF*_) does better than all other models for these mice. (C) We further investigate the difference between Age and vFI predictors in terms of feature importance. Features lying along the diagonal are important for both Age and vFI predictions. (D) Predicting FI score from video features extracted from videos of shorter durations. We used video features generated from videos with shorter durations (first 5 and 20 minutes) to investigate the loss in accuracy in predicting age and FI score. We used the random forest model trained with features generated from 60-minute videos as a baseline model for comparison. We found a diminished loss in accuracy using shorter videos. (E) To see how much training data is realistically needed, we performed a simulation study where we allocated different percentage of total data to training. As expected, there is a general downward (upward) trend in MAE, RMSE (R^2^ with an increasing percentage of data allocated to training set. Indeed, a smaller training set (*<* 80% training) can reach a similar training performance.

